# Re-oxidation of cytosolic NADH is a major contributor to the high oxygen requirements of the thermotolerant yeast *Ogataea parapolymorpha* in oxygen-limited cultures

**DOI:** 10.1101/2021.04.30.442227

**Authors:** Wijbrand J. C. Dekker, Hannes Juergens, Raúl A. Ortiz-Merino, Christiaan Mooiman, Remon van den Berg, Astrid Kaljouw, Robert Mans, Jack T. Pronk

## Abstract

Thermotolerance is an attractive feature for yeast-based industrial ethanol production. However, incompletely understood oxygen requirements of known thermotolerant yeasts are incompatible with process requirements. To study the magnitude and molecular basis of these oxygen requirements in the facultatively fermentative, thermotolerant yeast *Ogataea parapolymorpha*, chemostat studies were performed under defined oxygen-sufficient and oxygen-limited cultivation regimes. The minimum oxygen requirements of *O. parapolymorpha* were found to be at least an order of magnitude larger than those of the thermotolerant yeast *Kluyveromyces marxianus*. This high oxygen requirement coincided with absence of glycerol formation, which plays a key role in NADH reoxidation in oxygen-limited cultures of other facultatively fermentative yeasts. Co-feeding of acetoin, whose reduction to 2,3-butanediol can reoxidize cytosolic NADH, supported a 2.5-fold higher biomass concentration in oxygen-limited cultures. The apparent inability of *O. parapolymorpha* to produce glycerol correlated with absence of orthologs of the *S. cerevisiae* genes encoding glycerol-3P phosphatase (Sc*GPP1*, Sc*GPP2*). Glycerol production was observed in aerobic batch cultures of a strain in which genes including key enzymes in mitochondrial reoxidation of NADH were deleted. However, transcriptome analysis did not identify a clear candidate for the responsible phosphatase. Expression of Sc*GPD2*, encoding NAD^+^-dependent glycerol-3P dehydrogenase, and Sc*GPP1* in *O. parapolymorpha* resulted in increased glycerol production in oxygen-limited chemostats, but glycerol production rates remained substantially lower than observed in *S. cerevisiae* and *K. marxianus*. These results identify a dependency on aerobic respiration for reoxidation of NADH generated in biosynthesis as a key factor in the unexpectedly high oxygen requirements of *O. parapolymorpha*.

**Importance:** Thermotolerant yeasts hold great potential for anaerobic fermentation processes but their application is so far hampered by incompletely understood oxygen requirements. Based on quantitative physiological studies in oxygen-limited chemostat cultures, this study shows that the thermotolerant yeast *Ogataea parapolymorpha* has a much higher oxygen requirement than other, previously investigated facultatively fermentative yeasts. The large oxygen requirement of *O. parapolymorpha* was linked to an inability of oxygen-limited cultures to efficiently reoxidize NADH formed in biosynthetic processes by glycerol formation. These results provide a basis for reducing the oxygen requirements of *O. parapolymorpha* by targeted metabolic engineering. In addition, this study shows that diversity of oxygen requirements should be taken into account in selecting yeast species for application in anaerobic or oxygen-limited industrial processes.

## Introduction

Microbial biotechnology offers promising options for replacing petrochemically produced chemicals by sustainable bio-based alternatives (1, 2). For example, microbial production of ethanol as a renewable transport fuel, based on plant carbohydrates as renewable feedstocks, is already applied on a large scale. The current annual production volume of 87 Mton ethanol (3) is almost exclusively produced with the yeast *Saccharomyces cerevisiae* (4, 5). In well-established ‘first-generation’ ethanol processes, this yeast converts readily fermentable sugars, derived from corn starch or sugar cane, at high rates, titers and yields (6). Use of engineered pentose-fermenting *S. cerevisiae* strains for conversion of pentose-containing lignocellulosic hydrolysates, generated from agricultural residues such as corn stover and sugar cane bagasse, is currently explored at industrial scale (4).

Economic viability of the microbial production of ethanol and other large-product-volume fermentation products requires near-theoretical product yields on carbohydrate feedstocks. These can only be obtained in the absence of respiration, and with minimum processing costs (4, 5). In practice, industrial ethanol fermentation is performed in large tanks, which readily become and remain anaerobic as a consequence of vigorous carbon dioxide production by fermenting yeast cells. The popularity of *S. cerevisiae* in these processes is related to its high fermentation rates under anaerobic conditions, its high innate ethanol tolerance, its tolerance to low pH and increasingly also to its amenability to modern genome-editing techniques (7–10).

*S. cerevisiae* is a mesophilic yeast, which grows optimally at temperatures of approximately 35 °C (11). Ethanol fermentation at higher temperatures could reduce cooling costs of large-scale bioreactors and, moreover, enable improved integration of plant-polysaccharide hydrolysis with fermentation of the released monomeric sugars. Integration of enzymatic polysaccharide hydrolysis and fermentation (simultaneous saccharification and fermentation, SSF (12)) in a single unit operation could simplify processing and prevent feedback inhibition of polysaccharide hydrolysis by released monosaccharides (12, 13). Such an integration is especially relevant for second-generation bioethanol production, in which physically and/or chemically pre-treated lignocellulosic plant biomass is hydrolyzed to monomeric sugars by fungal hydrolases, which typically operate at temperatures of 50 – 80 °C (14, 15). However, the efficiency of yeast-based SSF processes is currently constrained by the limited temperature range of *S. cerevisiae*.

Since supra-optimal temperatures potentially affect all proteins in a cell (16), substantial extension of the temperature range of *S. cerevisiae* by metabolic engineering may prove to be an elusive target. Indeed, elegant adaptive laboratory evolution and metabolic engineering studies aimed at improving the thermotolerance of *S. cerevisiae*, e.g. by engineering its sterol composition, have only enabled modest improvements of its maximum growth temperature (17–19). Exploration of naturally thermotolerant, facultatively fermentative yeasts such as *Ogataea* sp. (*Hansenula* sp.)(20) and *Kluyveromyces marxianus*, whose temperature maxima can reach 50 °C (21, 22), appears to offer an attractive alternative approach. As is observed for the large majority of yeast species, these thermotolerant yeasts readily ferment sugars to ethanol under oxygen-limited conditions (23–25). However, with few exceptions, Saccharomycotina yeasts such as *O. parapolymorpha* and *K. marxianus*, whose evolutionary history did not involve the whole-genome duplication event (WGD) that shaped the genomes of *S. cerevisiae* and closely related species (26), cannot grow under strictly anaerobic conditions (23, 27). This inability is generally attributed to direct requirements for molecular oxygen and/or requirements for mitochondrial respiration in essential biosynthetic reactions and processes (28).

The molecular basis for the biosynthetic oxygen requirements has not been identified for all facultatively fermentative yeasts and can encompass processes such as sterol synthesis and/or sterol uptake, unsaturated fatty-acid synthesis, pyrimidine biosynthesis and synthesis of cofactors such as coenzyme A and pyridine nucleotides (28). In yeasts, molecular oxygen is required for unsaturated fatty acid synthesis due to involvement of the cytochrome-b5 Δ9-desaturase Ole1 (29). Similarly, involvement of multiple mono-oxygenases explains the requirement for synthesis of ergosterol (reviewed in (30)). Anaerobic cultures of *S. cerevisiae* are therefore routinely supplemented with ergosterol and Tween 80, an oleic acid ester that serves as source of unsaturated fatty acids (31). In most non-*Saccharomyces* yeasts, pyrimidine synthesis depends on a mitochondrial, respiration-coupled dihydro-orotate dehydrogenase (DHODase, Ura9). In contrast, *S. cerevisiae* only harbors a cytosolic DHODase (Ura1), which couples an oxygen- and respiration-independent dihydro-orotate oxidation to fumarate reduction (26, 32–34). *Kluyveromyces* sp. represent an evolutionary intermediate containing both cytosolic and mitochondrial DHODase variants, whose expression is regulated in response to oxygen availability (35). Development of metabolic engineering strategies for eliminating oxygen requirements of thermotolerant non-*Saccharomyces* yeasts requires elucidation of the underlying oxygen- and/or respiration-dependent biochemical reactions.

The goal of the present study was to investigate oxygen requirements of the thermotolerant yeast *O. parapolymorpha*. To this end, we first characterized its physiological and transcriptional responses to oxygen limitation in chemostat cultures and compared previously reported data (35) for *K. marxianus* and *S. cerevisiae*. Co-feeding of acetoin was tested to explore a possible impact of cytosolic NADH oxidation on the observed physiology of *O. parapolymorpha* under these conditions. Subsequently, we analyzed the genome of *O. parapolymorpha* for the presence or absence of orthologs of genes that have been implicated in the (in)ability of other yeasts to grow anaerobically and, in particular, in the ability to produce glycerol as ‘redox sink’ for reoxidation of NADH formed in biosynthetic reactions. Glycerol metabolism in *O. parapolymorpha* was further explored by studying growth of a mutant strain in which key genes involved in mitochondrial respiratory-chain linked NADH oxidation had been deleted. Based on the results of these analyses, we explored metabolic engineering of redox metabolism in *O. parapolymorpha* by expressing *S. cerevisiae* genes involved in glycerol production.

## Results

### Oxygen requirements of *O. parapolymorpha* in oxygen-limited chemostat cultures

To quantitatively assess biosynthetic oxygen requirements of *O. parapolymorpha*, physiological responses of the wild-type strain CBS11895 (DL-1)(36) were studied under two oxygenation regimes in chemostat cultures that were grown on glucose at a dilution rate of 0.10 h^-1^ (**Fig. 1**). Physiological parameters of *O. parapolymorpha* were compared to results obtained under the same cultivation conditions in a recent study on *S. cerevisiae* CEN.PK113-7D and *K. marxianus* CBS6556 (35).

**Figure 1.**
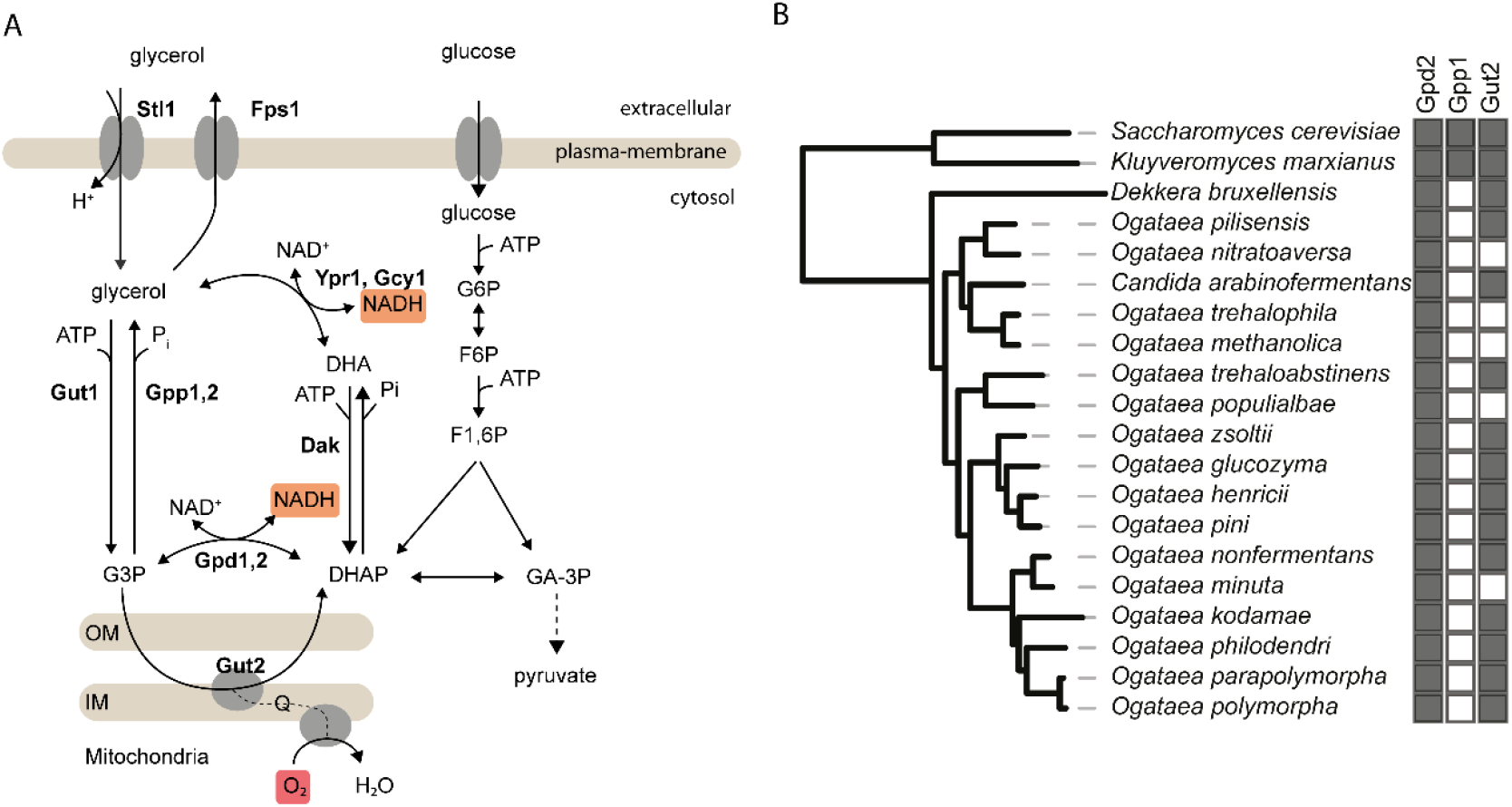
(**A**) Possible key reactions in glycerol metabolism and upper glycolysis in yeast. (**B**) Distribution of *S. cerevisiae* orthologs of structural genes encoding glycerol-3P dehydrogenase (Gpd2), glycerol-3P phosphatase (Gpp1) and FAD-dependent mitochondrial glycerol-3P dehydrogenase (Gut2) in *Ogataea* sp., *Brettanomyces* (syn. *Dekkera bruxellensis*), *K. marxianus* and *S. cerevisiae*. Data represents homology search results with *S. cerevisiae* S288C proteins as queries against whole genome translated sequences, present (black) and absent (white) orthologs are indicated. Strains are mapped to the phylogenetic tree of *Saccharomycotina* yeasts (48).

In fully aerobic chemostat cultures sparged with air (0.5 L min^-1^), growth of *O. parapolymorpha* was glucose limited and sugar dissimilation occurred exclusively via respiration, as indicated by a respiratory quotient (RQ) close to 1 (**Table 1**). The apparent biomass yield on glucose in these cultures was approximately 10 % higher than previously reported (37) due to co-consumption of ethanol, which was used as solvent for the anaerobic growth factor ergosterol. When oxygen availability was decreased by sparging with a mixture of N_2_ and air (0.5 L min^-1^, oxygen content 840 ppm), the apparent biomass yield on glucose in steady-state cultures was approximately four-fold lower than in fully aerobic cultures (0.15 g g^-1^ and 0.59 g g^-1^, respectively, **Table 1**). Moreover, a high residual glucose concentration (15.9 g L^-1^) indicated that growth was limited by oxygen rather than by glucose. Respiro-fermentative glucose metabolism in the oxygen-limited cultures was evident from a respiratory quotient (RQ) of 10.7 and a specific ethanol-dissimilation rate of 4.8 mmol (g biomass)^-1^ h^-1^. In contrast to results obtained in the same cultivation system with *K. marxianus* and *S. cerevisiae* (35), reduction of the O_2_ content of the inlet gas to < 0.5 ppm O_2_ led to complete wash-out of the *O. parapolymorpha* chemostat cultures.

**Table 1.** Physiology of yeast strains in glucose-grown chemostat cultures under different aeration regimes. Cultures were grown at pH 6.0 on synthetic medium with urea as nitrogen source, with glucose concentrations in the feed of aerobic and oxygen-limited cultures of 7.5 and 20 g L^-1^; respectively. Data for *K. marxianus* and *S. cerevisiae* were obtained from a previous study performed under identical conditions (35). Media were supplemented with the anaerobic growth factors ergosterol and Tween 80, except for aerobic cultures of *O. parapolymorpha*, from which Tween 80 was omitted to prevent excessive foaming. Data are represented as average ± SEM of data obtained from independent chemostat cultures of each strain. Ethanol concentrations were corrected for evaporation (35). Negative and positive biomass-specific conversion rates (q) represent consumption and production rates, respectively; subscript x denotes biomass dry weight. Calculations not applicable due to co-consumption of ethanol (-) and concentrations below detection limit (B.D., concentrations < 0.01 mM) are indicated.

| Yeast strain | *O. parapolymorpha*<br>CBS11895 |  | *K. marxianus*<br>CBS6556 |  | *S. cerevisiae*<br>CEN.PK113-7D |  |
| --- | --- | --- | --- | --- | --- | --- |
| O<sub>2</sub> in inlet gas (ppm) | 21·10<sup>4</sup> | 840 | 21·10<sup>4</sup> | 840 | 21·10<sup>4</sup> | 840 |
| Replicates | 2 | 2 | 2 | 5 | 3 | 3 |
| Biomass (g<sub>x</sub> L<sup>-1</sup>) | 4.33 ± 0.04 | 0.62 ± 0.01 | 3.79 ± 0.02 | 1.57 ± 0.10 | 4.22 ± 0.06 | 2.29 ± 0.04 |
| Residual glucose (g L<sup>-1</sup>) | B.D. | 15.92 ± 0.01 | B.D. | 0.10 ± 0.03 | B.D. | 0.07 ± 0.00 |
| **Biomass-specific rates** |  |  |  |  |  |  |
| Specific growth rate (h<sup>-1</sup>) | 0.10 ± 0.00 | 0.10 ± 0.00 | 0.10 ± 0.00 | 0.11 ± 0.01 | 0.10 ± 0.00 | 0.10 ± 0.00 |
| q<sub>glucose</sub> (mmol g<sub>x</sub><sup>-1</sup> h<sup>-1</sup>) | -0.94 ± 0.02 | -3.67 ± 0.14 | -1.05 ± 0.00 | -7.46 ± 0.30 | -0.95 ± 0.03 | -4.59 ± 0.10 |
| q<sub>ethanol</sub> (mmol g<sub>x</sub><sup>-1</sup> h<sup>-1</sup>) | -0.48 ± 0.04 | 4.75 ± 0.23 | -0.52 ± 0.00 | 11.49 ± 0.44 | -0.44 ± 0.03 | 7.48 ± 0.10 |
| q<sub>glycerol</sub> (mmol g<sub>x</sub><sup>-1</sup> h<sup>-1</sup>) | B.D. | 0.02 ± 0.00 | B.D. | 1.12 ± 0.06 | B.D. | 0.45 ± 0.01 |
| q<sub>succinate</sub> (mmol g<sub>x</sub><sup>-1</sup> h<sup>-1</sup>) | B.D. | 0.00 ± 0.00 | B.D. | 0.02 ± 0.00 | B.D. | 0.00 ± 0.00 |
| q<sub>O<sub>2</sub></sub> (mmol g<sub>x</sub><sup>-1</sup> h<sup>-1</sup>) | -2.35 ± 0.08 | -0.60 ± 0.01 | -3.52 ± 0.05 | -0.23 ± 0.02 | -2.61 ± 0.11 | -0.15 ± 0.01 |
| q<sub>CO<sub>2</sub></sub> (mmol g<sub>x</sub><sup>-1</sup> h<sup>-1</sup>) | 2.89 ± 0.12 | 6.42 ± 0.22 | 3.73 ± 0.03 | 10.5 ± 0.5 | 2.82 ± 0.10 | 7.91 ± 0.53 |
| **Stoichiometries** |  |  |  |  |  |  |
| RQ (qCO<sub>2</sub>/qO<sub>2</sub>) | 1.23 ± 0.09 | 10.7 ± 0.2 | 1.06 ± 0.01 | 49.3 ± 7.5 | 1.08 ± 0.02 | 52.2 ± 2.4 |
| Y<sub>glycerol/x</sub> (mmol g<sub>x</sub><sup>-1</sup>) | B.D. | 0.24 ± 0.05 | B.D. | 10.7 ± 0.7 | B.D. | 4.66 ± 0.08 |
| Y<sub>X/glucose</sub> (g<sub>x</sub> g<sup>-1</sup>) | 0.59 ± 0.00 | 0.15 ± 0.01 | 0.53 ± 0.00 | 0.08 ± 0.01 | 0.57 ± 0.01 | 0.12 ± 0.00 |
| Y<sub>ethanol/glucose</sub> (g g<sup>-1</sup>) | - | 0.33 ± 0.03 | - | 0.39 ± 0.00 | - | 0.42 ± 0.02 |
| Y<sub>X/O<sub>2</sub></sub> (g<sub>x</sub> mmol<sup>-1</sup>) | 0.04 ± 0.00 | 0.17 ± 0.00 | 0.03 ± 0.01 | 0.50 ± 0.08 | 0.04 ± 0.00 | 0.64 ± 0.03 |
| **Recoveries (out/in)** |  |  |  |  |  |  |
| Carbon (%) | 100.3 ± 0.4 | 94.4 ± 4.4 | 100.5 ± 0.1 | 91.1 ± 2.0 | 99.9 ± 0.7 | 101.2 ± 3.3 |
| Degree of reduction(%) | 97.8 ± 0.5 | 92.0 ± 4.3 | 98.8 ± 0.1 | 98.4 ± 0.7 | 98.4 ± 0.7 | 100.1 ± 0.8 |

*S. cerevisiae* can grow anaerobically in media supplemented with B-type vitamins, sterols and a source of UFA (31, 38, 39). In *K. marxianus*, which lacks a functional sterol-uptake system, sterol synthesis was recently identified as the key oxygen-requiring process during growth in such media (35). Based on UFA and sterol contents of aerobically grown *S. cerevisiae* biomass, the required oxygen uptake rate for synthesis of these lipids at a specific growth rate of 0.10 h^-1^ corresponds to approximately 0.01 mmol O_2_ (g biomass)^-1^ h^-1^ (39). This requirement is 60-fold lower than the biomass-specific oxygen consumption rate of 0.60 mmol O_2_ (g biomass)^-1^ h^-1^ observed in oxygen-limited cultures of *O. parapolymorpha*.

A comparison of oxygen-uptake and ethanol-production rates indicated that, in the oxygen-limited *O. parapolymorpha* cultures, approximately 3 % of the total consumed glucose by the oxygen-limited cultures was respired. In contrast, under the same oxygen-limited regime, chemostat cultures of *S. cerevisiae* and *K. marxianus* showed specific oxygen-consumption rates below 0.25 mmol (g biomass)^-1^ h^-1^, RQ values above 50 and very low residual glucose concentrations (**Table 1**)(35). For these two yeasts, the fraction of the total consumed glucose that was respired amounted to only approximately 0.5 %.

In comparison with oxygen-limited cultures of *K. marxianus* and *S. cerevisiae* (**Table 1**), those of *O. parapolymorpha* showed an over ten-fold lower biomass-specific rate of glycerol production. Glycerol formation plays an important role in anaerobic and severely oxygen-limited cultures of several yeasts by reoxidizing a surplus of cytosolic NADH formed in biosynthetic reactions (40–43). In fully anaerobic cultures of *S. cerevisiae* and in severely oxygen-limited cultures of *K. marxianus*, glycerol formation has been reported to be 7-12 mmol (gram biomass)^-1^(35). At a dilution rate of 0.10 h^-1^, reoxidation of an equivalent amount of NADH by aerobic respiration would require an uptake rate of 0.4 to 0.6 mmol O_2_ g (biomass)^-1^ h^-1^, which closely corresponds with the observed oxygen consumption rate of the oxygen-limited *O. parapolymorpha* cultures.

In some yeast species, an insufficient capacity for glycerol production causes an inability to grow under severe oxygen limitation. This phenomenon, which is referred to as the ‘Custers effect’, can be complemented by supplementation of oxygen-limited cultures with acetoin, whose NADH-dependent reduction to 2,3-butanediol can replace glycerol formation as a cellular ‘redox sink’ (40, 44). Addition of acetoin to oxygen-limited chemostat cultures of *O. parapolymorpha* led to an increase of the steady-state biomass concentration from 0.62 to 1.57 g (biomass)^-1^ L^-1^ and metabolism became more fermentative, as indicated by a higher rate of ethanol production and higher RQ (**Table 3**).

**Table 2.** Physiology of *O. parapolymorpha* wild-type and mutant strains in glucose-grown aerobic batch and chemostat cultures. Aerobic chemostat cultures were grown on synthetic medium with and 20 g L^-1^ glucose as carbon- and energy substrate. Batch cultures were grown on synthetic medium with a reduced 7.5 g L^-1^ glucose concentration. Data are represented as average ± SEM of data obtained from independent chemostat cultures of each strain. Ethanol concentrations were corrected for evaporation. Positive and negative biomass-specific conversion rates (q) represent consumption and production rates, respectively; subscript x denotes biomass dry weight. N.D.: not determined values; and B.D.: compounds below detection limit (concentrations < 0.01 mM).

| *O. parapolymorpha* strain | CBS11895 | IMX2167 | CBS11895 | IMX2167 |
| --- | --- | --- | --- | --- |
| Relevant genotype | wt | *ndh1-3Δ gut2Δ* | wt | *ndh1-3Δ gut2Δ* |
| Replicates | 2 | 2 | 2 | 2 |
| **Biomass-specific rates** |  |  |  |  |
| Specific growth rate (h<sup>-1</sup>) | 0.36 ± 0.006 | 0.26 ± 0.00 | 0.10 ± 0.00 | 0.10 ± 0.00 |
| q<sub>glucose</sub> (mmol g<sub>x</sub> h<sup>-1</sup>) | -3.90 ± 0.15 | -2.73 ± 0.06 | -1.08 ± 0.03 | -1.04 ± 0.00 |
| q<sub>ethanol</sub> (mmol g<sub>x</sub> h<sup>-1</sup>) | B.D. | 0.22 ± 0.08 | B.D. | B.D. |
| q<sub>glycerol</sub> (mmol g<sub>x</sub> h<sup>-1</sup>) | B.D. | 0.34 ± 0.01 | B.D. | B.D. |
| q<sub>O2</sub> (mmol g<sub>x</sub> h<sup>-1</sup>) | N.D. | -3.65 ± 0.05 | -2.69 ± 0.05 | -2.14 ± 0.00 |
| q<sub>CO2</sub> (mmol g<sub>x</sub> h<sup>-1</sup>) | N.D. | 4.34 ± 0.01 | 2.82 ± 0.03 | 2.25 ± 0.00 |
| **Stoichiometries** |  |  |  |  |
| RQ (qCO<sub>2</sub>/qO<sub>2</sub>) | N.D. | 1.19 ± 0.02 | 1.05 ± 0.01 | 1.05 ± 0.00 |
| Y<sub>X/glucose</sub> (g<sub>x</sub> g<sup>-1</sup>) | 0.52 ± 0.01 | 0.51 ± 0.01 | 0.51 ± 0.00 | 0.52 ± 0.00 |
| Y<sub>X/O2</sub> (g<sub>x</sub> mmol<sup>-1</sup>) | N.D. | 0.07 ± 0.00 | 0.04 ± 0.00 | 0.05 ± 0.00 |
| **Recoveries (out/in)** |  |  |  |  |
| Carbon (%) | N.D. | 97.8 ± 1.0 | 99.3 ± 1.2 | 98.7 ± 0.3 |

**Table 3.** Physiology of *O. parapolymorpha* strains expressing *S. cerevisiae* glycerol pathway genes in glucose-grown oxygen-limited chemostat cultures. Cultures were grown at pH 6.0 on synthetic medium with urea as nitrogen source and 20 g L^-1^ glucose as carbon and energy substrate, media contained anaerobic growth factors ergosterol and Tween 80. Data are represented as average ± SEM of data obtained from independent chemostat cultures of each strain. Ethanol concentrations were corrected for evaporation (35). Positive and negative biomass-specific conversion rates (q) represent consumption and production rates, respectively; subscript x denotes biomass dry weight. B.D.: concentrations below detection limit (concentration < 0.01 mM).

| Yeast strain | CBS11895 | IMX2119 | IMX2587 | IMX2588 | CBS11895 |
| --- | --- | --- | --- | --- | --- |
| Relevant genotype | wt | *ScGPP1* | *ScGPD2* | *ScGPP1*<br>*ScGPD2* | wt |
| O<sub>2</sub> in inlet gas (ppm) | 840 | 840 | 840 | 840 | 840 |
| Acetoin | - | - | - | - | yes |
| Replicates | 2 | 3 | 2 | 3 | 3 |
| Biomass (g<sub>x</sub> L<sup>-1</sup>) | 0.62 ± 0.01 | 0.91 ± 0.03 | 0.66 ± 0.06 | 0.86 ± 0.05 | 1.57 ± 0.10 |
| Residual glucose (g L<sup>-1</sup>) | 15.92 ± 0.01 | 14.33 ± 0.35 | 15.71 ± 0.40 | 14.98 ± 0.37 | 9.41 ± 0.24 |
| **Biomass-specific rates** |  |  |  |  |  |
| Specific growth rate (h<sup>-1</sup>) | 0.10 ± 0.00 | 0.11 ± 0.01 | 0.10 ± 0.01 | 0.12 ± 0.01 | 0.11 ± 0.00 |
| q<sub>glucose</sub> (mmol g<sub>x</sub><sup>-1</sup> h<sup>-1</sup>) | -3.67 ± 0.14 | -3.69 ± 0.25 | -2.97 ± 0.12 | -3.92 ± 0.28 | -4.11 ± 0.07 |
| q<sub>ethanol</sub> (mmol g<sub>x</sub><sup>-1</sup> h<sup>-1</sup>) | 4.75 ± 0.23 | 4.92 ± 0.30 | 4.72 ± 0.14 | 7.26 ± 0.44 | 5.90 ± 0.23 |
| q<sub>glycerol</sub> (mmol g<sub>x</sub><sup>-1</sup> h<sup>-1</sup>) | 0.02 ± 0.00 | 0.18 ± 0.01 | 0.03 ± 0.01 | 0.22 ± 0.00 | 0.03 ± 0.00 |
| q<sub>succinate</sub> (mmol g<sub>x</sub><sup>-1</sup> h<sup>-1</sup>) | B.D. | B.D. | 0.02 ± 0.02 | 0.01 ± 0.01 | 0.02 ± 0.00 |
| q<sub>acetoin</sub> (mmol g<sub>x</sub><sup>-1</sup> h<sup>-1</sup>) | - | - | - | - | -7.45 ± 0.18 |
| q<sub>O<sub>2</sub></sub> (mmol g<sub>x</sub><sup>-1</sup> h<sup>-1</sup>) | -0.60 ± 0.01 | -0.44 ± 0.02 | -0.59 ± 0.04 | -0.37 ± 0.02 | -0.29 ± 0.01 |
| q<sub>CO<sub>2</sub></sub> (mmol g<sub>x</sub><sup>-1</sup> h<sup>-1</sup>) | 6.42 ± 0.22 | 6.41 ± 0.46 | 5.33 ± 0.26 | 6.45 ± 0.24 | 7.00 ± 0.24 |
| **Stoichiometries** |  |  |  |  |  |
| RQ (qCO<sub>2</sub>/qO<sub>2</sub>) | 10.7 ± 0.2 | 14.4 ± 0.5 | 9.11 ± 0.16 | 17.4 ± 1.0 | 24.2 ± 0.5 |
| Y<sub>glycerol/x</sub> (mmol g<sub>x</sub><sup>-1</sup>) | 0.24 ± 0.05 | 1.71 ± 0.05 | 0.30 ± 0.09 | 1.93 ± 0.15 | 0.32 ± 0.02 |
| Y<sub>X/glucose</sub> (g<sub>x</sub> g<sup>-1</sup>) | 0.15 ± 0.01 | 0.16 ± 0.00 | 0.21 ± 0.01 | 0.16 ± 0.01 | 0.14 ± 0.00 |
| Y<sub>ethanol/glucose</sub> (g g<sup>-1</sup>) | 0.33 ± 0.03 | 0.34 ± 0.01 | 0.41 ± 0.00 | 0.48 ± 0.03 | 0.37 ± 0.01 |
| $Y_{x/O_2}$ (g<sub>x</sub> mmol<sup>-1</sup>) | 0.17 ± 0.00 | 0.24 ± 0.00 | 0.18 ± 0.02 | 0.32 ± 0.03 | 0.37 ± 0.00 |
| **Recoveries (out/in)** |  |  |  |  |  |
| Carbon (%) | 94.4 ± 4.4 | 98.4 ± 1.3 | 102.8 ± 1.2 | 106.1 ± 3.3 | 96.1 ± 1.0 |
| Degree of reduction (%) | 92.0 ± 4.3 | 93.5 ± 1.8 | 102.9 ± 0.3 | 109.6 ± 3.3 | 99.5 ± 1.0 |

### Absence of orthologs of *S. cerevisiae* NAD^+^-dependent glycerol-3P phosphatase in *Ogataea* species

To study the molecular basis for the apparent absence of a functional glycerol pathway in the oxygen-limited *O. parapolymorpha* cultures, we investigated the presence of orthologs of *S. cerevisiae GPD1/2* and *GPP1/2*. These genes encode isoenzymes catalyzing the two key reactions of the *S. cerevisiae* glycerol pathway, NAD^+^-dependent glycerol-3P dehydrogenase and glycerol-3P phosphatase, respectively (45–47). A homology search in translated whole genome sequences of 16 strains of *Ogataea* species (48) revealed clear orthologs of Gpd, but no orthologs of Gpp were detected (**Fig. 1**). In this respect, *Ogataea* yeasts resembled the phylogenetically related genus *Brettanomyces* (syn. *Dekkera*) (**Fig. 1**) for which the occurrence of a Custers effect is well documented (49, 50). In the absence of an active glycerol-3P phosphatase, NAD^+^-dependent glycerol-3P dehydrogenase can still be involved in glycerolipid synthesis (51) and in the glycerol-3P shuttle, in which glycerol-3P is reoxidized to dihydroxyacetone phosphate by a mitochondrial, respiratory-chain-coupled glycerol-3P dehydrogenase (Gut2 in *S. cerevisiae*, (52–54), **Fig. 1**).

### Transcriptional responses of *O. parapolymorpha* to oxygen limitation

To further investigate responses of *O. parapolymorpha* to oxygen limitation, transcriptome analyses were performed on chemostat cultures grown under the aerobic and micro-aerobic regimes described above. These transcriptome data were first used to refine the genome annotation of a *de novo* assembled genome sequence of *O. parapolymorpha* CBS11895 obtained from long-read sequence data (**Data availability**). Data analysis focused on *O. parapolymorpha* genes with orthologs in *S. cerevisiae* and *K. marxianus* (**Fig. 2**). Transcriptional responses to oxygen limitation in *O. parapolymorpha* were compared with previously obtained transcriptome datasets (35, 55) of the latter two facultatively fermentative yeasts, grown under aerobic and oxygen-limited conditions. A global comparison indicated large differences in the transcriptional responses to these cultivation conditions (**Fig. 2**). The only shared global transcriptional responses of *O. parapolymorpha, K. marxianus*, and *S. cerevisiae* were a downregulation, in the oxygen-limited cultures, of genes involved in metabolism of alternative non-glucose carbon sources and in isocitrate metabolism. These responses are in line with an oxygen dependency for the dissimilation of these substrates and for a key role of the tricarboxylic acid cycle in respiratory glucose metabolism, respectively. At first glance, the different global transcriptional responses of these three yeasts appear to indicate a completely different wiring of their oxygen-responsive transcriptional regulation networks. However, it should be noted that high residual glucose concentrations in the oxygen-limited *O. parapolymorpha* cultures (**Table 1**) may have influenced transcriptional regulation (56).

**Figure 2.**
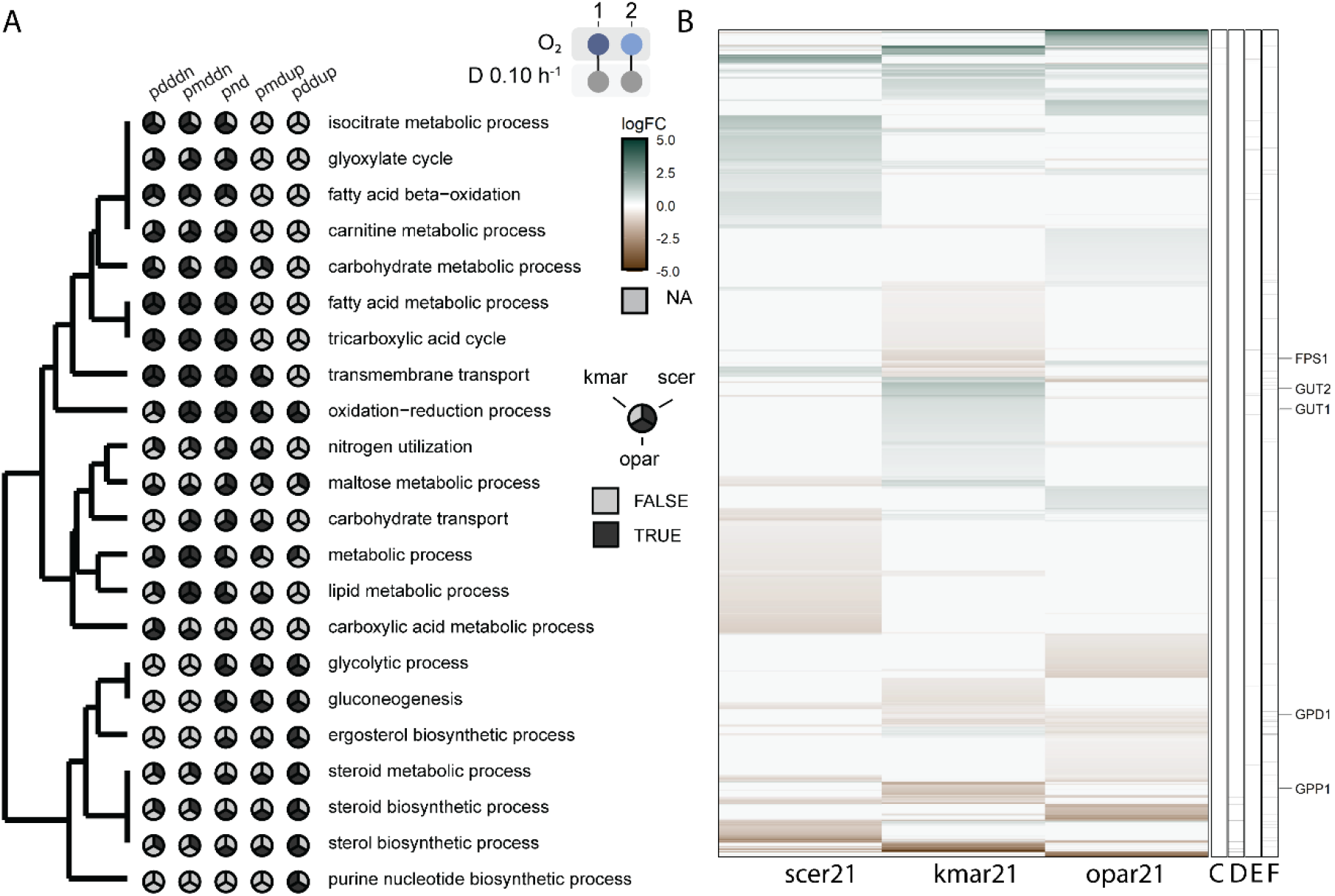
Genome-wide transcriptional responses of *O. parapolymorpha, K. marxianus* and *S. cerevisiae* to oxygen limitation. Chemostat cultures of *O. parapolymorpha* (opar) were grown at pH 6 on synthetic medium with urea as nitrogen source and either aerated with air (aerobic 21·10^4^ ppm O_2_, regime 1), or with a mixture of N_2_ and 840 ppm O_2_ (micro-aerobic, regime 2). Glucose concentrations in the feed of aerobic and oxygen-limited cultures were 7.5 and 20 g L^-1^; respectively. Data for *K. marxianus* (kmar) and *S. cerevisiae* (scer) were obtained from a previous study performed under identical conditions (35). (**A**) Comparison of gene-set enrichment analysis using GO-terms for biological processes in response to oxygen limitation (micro-aerobic vs. aerobic). Distinct directionalities were calculated with Piano (58) and indicated as distinct directional up (pddup), mixed directional up (pmdup), non-directional (pnd), mixed directional down (pmddn) and distinct directional down (pdddn). Hierarchical clustering was performed based on the frequency of enrichment of GO-terms in the three yeast strains. (**B)** Highlighted gene sets show similar log fold changes of orthologs in the three yeasts, (**C**) both in all three yeasts upregulated (**D**) both downregulated. (**E**) and (**F**) indicate *S. cerevisiae* genes that were found to be consistently (**E**) up-regulated or (**F**) down-regulated in anaerobic chemostat cultures across four nutrient limitation regimes (55).

In contrast to *S. cerevisiae*, which showed downregulation of genes associated with sterol metabolism, *O. parapolymorpha* and *K. marxianus* showed upregulation of genes associated with this GO-term (based on GO-term enrichment analysis). Sterol biosynthesis requires molecular oxygen and, under anaerobic or oxygen-limited conditions, *S. cerevisiae* can acquire ergosterol from the media. *K. marxianus* and several other pre-WGD yeast species lack a functional sterol import system (35, 57). This analysis of the transcriptome response of *O. parapolymorpha* suggested that this is also the case for this yeast.

Op*URA9*, which encodes the respiratory-chain-linked dihydroorotate dehydrogenase of *O. polymorpha*, showed an upregulation under oxygen-limited conditions, while Km*Ura9* was downregulated (**Fig. 3**). This response is consistent with the absence of a respiration-independent ScUra1/KmUra1-type dihydroorotate dehydrogenase in *O. parapolymorpha*.

**Figure 3.**
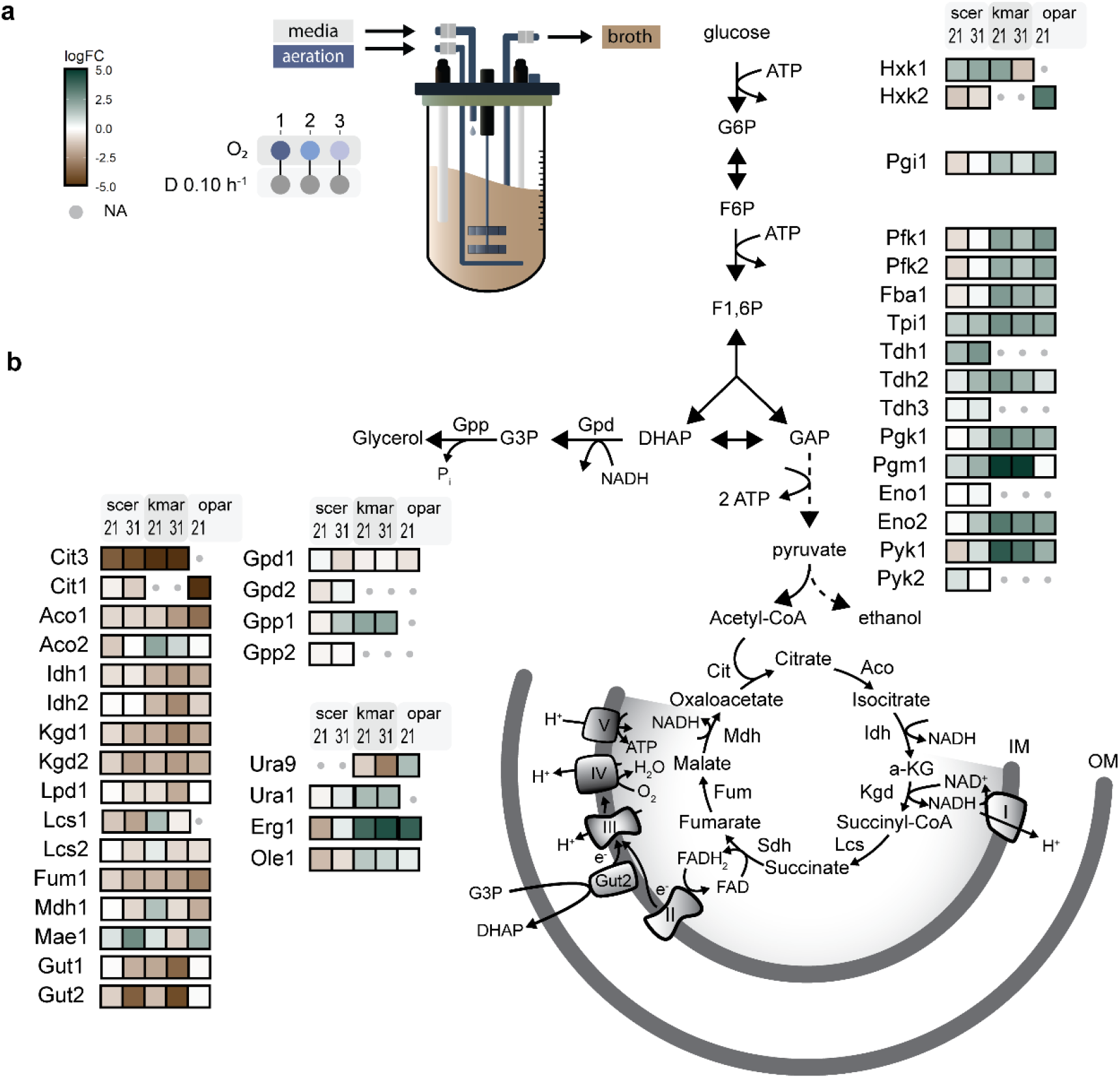
Transcriptional regulation of specific pathways and structural genes in *O. parapolymorpha, K. marxianus* and *S. cerevisiae* subjected to different aeration regimes. (**A**) Glucose-grown chemostat cultures (dilution rate 0.10 h^-1^) were aerated with air (regime 1), a gas mixture containing 840 ppm O_2_ (regime 2) or with nitrogen containing < 0.5 ppm O_2_ (regime 3) at a rate of 0.5 L min^-1^. *O. parapolymorpha* washed out under regime 3. RNAseq data are compared against aerobic, glucose-limited cultures (regime 1) as the reference. Transcriptome data for *K. marxianus* and *S. cerevisiae* were obtained from a previous study (35). (**B**) Biochemical reactions are represented by arrows and multiple reactions by dashed arrows. Boxes with colors indicate gene upregulation (blue-green) or downregulation (brown), intensities indicating the log 2 fold change (logFC) with color range capped to a maximum value of 5. Schematic representation of biochemical reactions and for ease of visualization not all reaction products or cofactors are depicted. Reactions are annotated with the corresponding *S. cerevisiae* ortholog name; absence of orthologs is indicated by grey dots. Abbreviations used; glucose-6-phosphate (G6P), fructose-6-phosphate (F6P), fructose-1,6-bisphosphate (F1,6P), dihydroxyacetone phosphate (DHAP), glycerol-3P (G3P), glyceraldehyde phosphate (GAP), inner-mitochondrial membrane (IM), outer-mitochondrial membrane (OM). Respiratory chain complexes are indicated by Roman numerals.

The importance of glycerol production for oxygen-limited growth of *S. cerevisiae* and *K. marxianus* was evident from a strong upregulation of *GPP1*, for which no ortholog was found in *O. parapolymorpha* (**Fig. 3**). This observation and the lack of a transcriptional response of the single *GPD* ortholog in *O. parapolymorpha* to oxygen limitation are in line with the conclusion that this yeast does not use glycerol formation as a redox sink during oxygen-limited growth.

### Glycerol production in aerobic cultures of an *O. parapolymorpha* strain lacking mitochondrial glycerol-3P dehydrogenase and alternative NADH dehydrogenases

The presence of orthologs of *S. cerevisiae GPD1/2* and *GUT2* in *O. parapolymorpha* suggested possible involvement of a glycerol-3P shuttle (52) to respiratory oxidation of the cytosolic NADH derived from glycolysis and from biosynthetic reactions. To further investigate redox metabolism in *O. parapolymorpha*, we studied growth and product formation in strain IMX2167. In this *O. parapolymorpha* strain, Op*GUT2* as well as the genes encoding the three cytosol-and matrix-facing alternative mitochondrial NADH dehydrogenases were deleted (59).

In aerobic chemostat cultures grown at a dilution rate of 0.10 h^-1^, conversion rates of strain IMX2167 did not substantially differ from those of the wild-type strain CBS11895 (**Table 2**). Apparently, as previously observed in aerobic cultures of corresponding mutant strains of *S. cerevisiae* (43), mechanisms such as an ethanol-acetaldehyde shuttle could compensate for the absence of mitochondrial glycerol-3P dehydrogenase and external NADH dehydrogenases. In strain IMX2167, in which also internal alternative NADH dehydrogenase was absent, such a shuttle mechanism could then couple reoxidation of cytosolic NAHD to the *O. parapolymorpha* complex-I NADH dehydrogenase (43).

In contrast to the absence of a clear phenotype in the chemostat cultures, aerobic batch cultures of strain IMX2167 showed a lower specific growth rate than the wild-type strain CBS11895 (0.26 h^-1^ and 0.36 h^-1^, respectively, **Table 2**). Glycerol production suggested that, despite the absence of an Sc*GPP1* ortholog, *O. parapolymorpha* contains a phosphatase that can use glycerol-3P as a substrate. In the genome sequence of strain CBS11895, 24 genes were annotated with the GO-term ‘phosphatase activity’ (GO:0016791). However, although all these genes were transcribed (log Counts per million (CPM) > 3.5), none showed log FC > 2 between the wild-type and strain IMX2167 in either aerobic batch cultures or aerobic chemostat cultures (**Table 2**).

### Metabolic engineering of glycerol metabolism in *O. parapolymorpha*

Based on analysis of the *O. parapolymorpha* genome (**Fig. 1**), we investigated whether expression of the *S. cerevisiae* glycerol-3P phosphatase ScGpp1 in *O. parapolymorpha* supported glycerol production by oxygen-limited cultures of *O. parapolymorpha*. To this end, an expression cassette in which the coding region of *ScGPP1* was expressed from the Op*PMA1* promoter was integrated into the genome of *O. parapolymorpha* CBS11895. In oxygen-limited chemostat cultures, the resulting engineered strain IMX2119 showed a 9-fold higher biomass-specific rate of glycerol formation than observed in the wild-type strain (0.18 mmol (g biomass)^-1^ h^-1^ and 0.02 mmol (g biomass)^-1^ h^-1^, respectively, **Table 3, Fig. 4A**). A further increase of the glycerol production rate to 0.22 mmol (g biomass)^-1^ h^-1^ was observed when, in addition to the expression cassette for Sc*GPP1*, a second expression cassette was integrated in which Sc*GPD2* was expressed from the Op*TEF1* promoter (strain IMX2588, **Table 3, Fig. 4A**). Integration of only the Sc*GPD2* cassette (strain IMX2587) did not lead to a significantly higher rate of glycerol production in oxygen-limited cultures than observed in the wild-type strain (**Table 3**).

**Figure 4.**
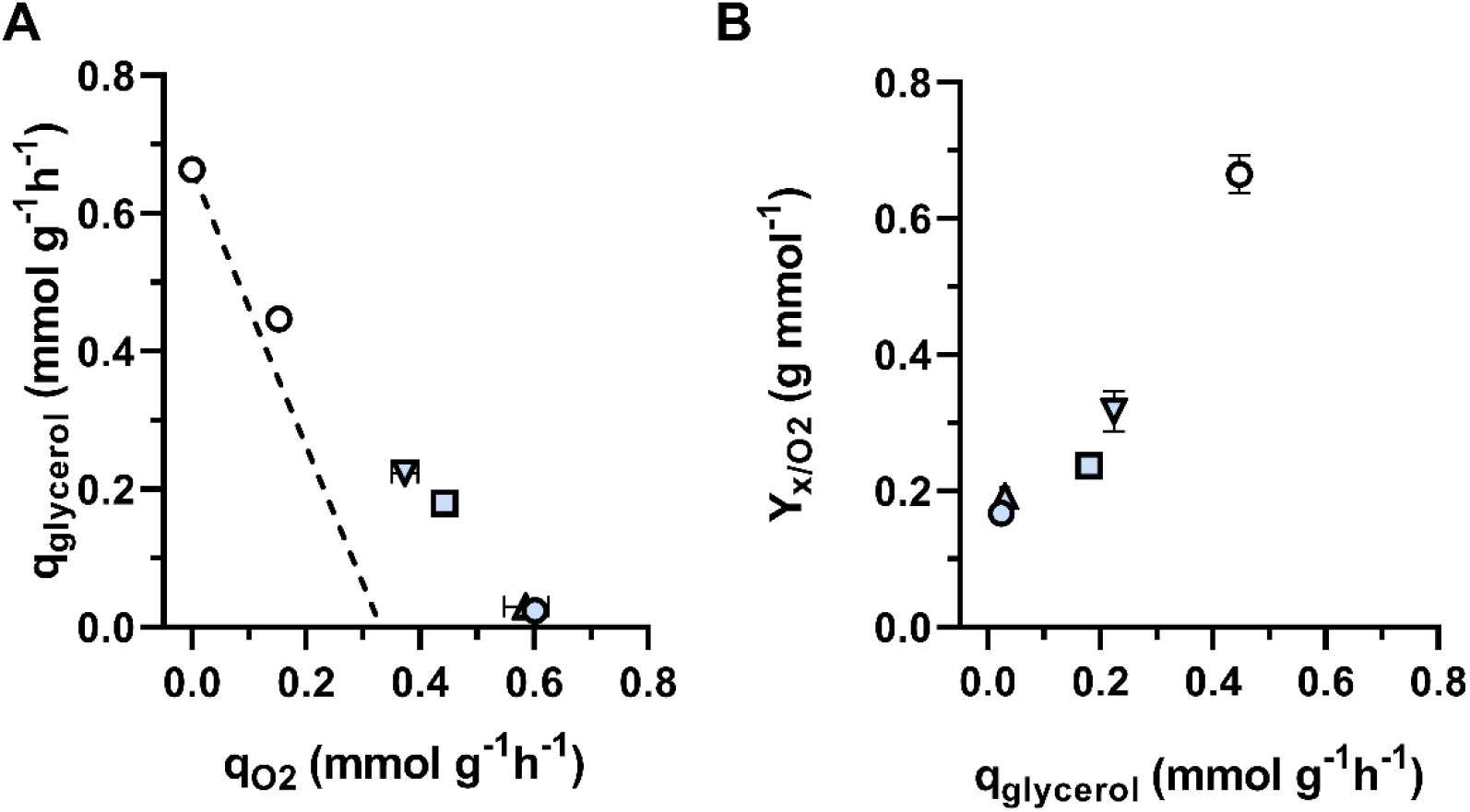
Impact of heterologous expression of Sc*GPP1* and/or Sc*GPD2* expression in *O. parapolymorpha* CBS11895 on glycerol production in oxygen-limited chemostat cultures. Biomass specific conversion rates (q) of *O. parapolymorpha* strains were measured in oxygen-limited conditions (dilution rate 0.10 h^-1^), aerated at 500 mL min^-1^ with a gas mixture containing 840 ppm O_2_. Data of anaerobic and oxygen-limited *S. cerevisiae* cultures, grown in the same experimental set-up where derived from (35). Symbols: white circle *S. cerevisiae* CEN.PK113-7D, blue circles *O. parapolymorpha* CBS11895, blue box IMX2119 (Sc*GPP1*), blue triangle up IMX2587 (Sc*GPD2*), blue triangle down IMX2588 (Sc*GPP1* Sc*GPD2*). Data are represented as average ± SEM of data obtained from independent chemostat cultures of each strain. (**A**) Biomass-specific glycerol production rate versus biomass-specific oxygen consumption rate. The dashed line depicts the stoichiometric relationship between glycerol and oxygen interpolated from the anaerobic rate of glycerol production by *S. cerevisiae* under the assumption that, for NADH reoxidation, consumption of one mol O_2_ is equivalent with the production of two mol glycerol. (**B**) Biomass yield on oxygen versus biomass-specific rate of glycerol production.

The higher biomass-specific rates of glycerol production by the Sc*GPP1* and Sc*GPP1*/Sc*GPD2* expressing *O. parapolymorpha* strains enabled higher biomass yields on oxygen under oxygen-limited conditions (0.24 and 0.32 g biomass mmol O_2_^-1^, respectively, versus 0.17 g biomass mmol O_2_^-1^ for the wild-type strain, **Table 3** and **Fig. 3B**). A shift towards a more fermentative metabolism was, in addition, evident from the RQ values of strains IMX2119 and IMX2588 (14.4 and 17.4, respectively), which were significantly higher than those of corresponding cultures of the wild-type strain (RQ of 10.7, **Table 3**).

In fully anaerobic chemostat cultures of *S. cerevisiae* CEN.PK113-7D grown at a dilution rate of 0.10 h^-1^, a biomass-specific rate of glycerol production of 0.67 mmol (g biomass)^-1^ h^-1^ has been reported (**Fig. 4A**)(35, 60). An even higher rate of glycerol production (1.1 mmol (g biomass)^-1^ h^-1^) has been reported for strains of another *S. cerevisiae* lineage grown under these conditions (41, 61). Assuming that biomass composition and biosynthetic pathways in *S. cerevisiae* CEN.PK113-7D and *O. parapolymorpha* CBS11895 lead to a similar net generation of NADH from biosynthetic reactions, the glycerol production rate of the Sc*GPP1*/Sc*GPD2* expressing *O*. parapolymorpha strain IMX2588 was approximately four-fold lower than required for reoxidation of all NADH generated in biosynthetic processes. A limiting capacity of the engineered glycerol pathway in oxygen-limited cultures of strain IMX2588 was also indicated by residual glucose concentrations, which were higher than in cultures of the wild-type strain CBS11895 supplemented with acetoin (**Table 3**).

## Discussion

The large majority of facultatively fermentative non-*Saccharomyces* yeasts require molecular oxygen and/or aerobic respiration for essential biosynthetic processes (23, 62). Comparative physiology studies across strains and species, as well as elucidation and differentiation of underlying biochemical mechanisms, require rigorous standardization of cultivation conditions to minimize oxygen contamination and differences in gas-transfer properties. In this study, chemostat cultivation under defined oxygenation regimes revealed surprisingly high oxygen requirements of oxygen-limited cultures of *Ogataea parapolymorpha* (previously *Hansenula polymorpha* (20)). These requirements far exceeded those reported for the facultatively fermentative pre-WGD yeasts *Kluyveromyces marxianus, K. lactis* and *Candida utilis* (*Cyberlindnera jadinii*)(41, 63). *O. parapolymorpha* is currently applied in aerobic industrial processes for production of heterologous proteins (64) and based on its thermotolerance and natural ability to metabolize D-xylose, is intensively investigated as a potential platform organism for second-generation ethanol production (21). In anaerobic industrial applications of *Saccharomyces* yeasts, such as beer fermentation, introduction of a brief aeration phase enables yeast cell to synthesize and intracellularly accumulate sterols and unsaturated fatty acids, which are then used during the subsequent anaerobic fermentation phase (65, 66). Since such a process intervention cannot address the large oxygen requirements of *O. parapolymorpha*, reduction or complete elimination of its oxygen requirements should be a priority metabolic-engineering target for development of industrial ethanol-producing strains.

Very low glycerol-production rates and an increased biomass yield upon acetoin co-feeding to oxygen-limited cultures identified reoxidation of NADH formed in biosynthetic reactions as a key contributor to the large oxygen requirements of *O. parapolymorpha*. A substantial oxygen requirement for fermentative growth (‘Custers effect’; (67)), absence of glycerol production and a stimulating effect of acetoin on oxygen-limited growth were previously observed in *Brettanomyces* (*Dekkera*) yeasts (40, 49, 67, 68). Further studies on *B. bruxellensis* related the Custers effect to absence of glycerol-3P phosphatase in cell extracts (49) and of an ortholog of the *S. cerevisiae GPP1/GPP2 genes* in its genome (69). The genera *Ogataea* and *Brettanomyces* both belong to the Pichiacaea family (70). Our observations on *O. parapolymorpha* and on the absence of Sc*GPP1*/Sc*GPP2* orthologs in *Ogataea* species, provide an incentive for further studies into occurrence of this intriguing phenomenon in Pichiacaea. In view of its fast growth in synthetic media (71) and its accessibility to modern genome-editing techniques (72, 73), *O. parapolymorpha* offers an interesting experimental platform to study the regulation networks and ecophysiological significance of the Custers effect. Formation of glycerol in aerobic cultures of a strain in which genes encoding key enzymes of respiratory NADH oxidation, including the mitochondrial flavoprotein NADH dehydrogenase (OpGut2), were deleted, raises the possibility that absence of glycerol formation in oxygen-limited cultures might be related to the operation of a glycerol-3P shuttle for mitochondrial oxidation of cytosolic NADH in this yeast (reviewed in (43)).

While heterologous expression of *S. cerevisiae GPD2* and *GPP1* in *O. parapolymorpha* increased glycerol production and biomass yield on oxygen in oxygen-limited cultures, it did not allow for complete fermentation of the supplied glucose. Increased expression of these heterologous genes, possibly combined with expression of a glycerol exporter and/or laboratory evolution under oxygen-limited conditions, is likely to be required for further improvement of oxygen-independent NADH reoxidation. Alternatively, use of other heterologous pathways in *O. parapolymorpha* that enable NADH-dependent reduction of acetyl-CoA to ethanol (74) or NADH oxidation via a pathway involving ribulose-1,5-bisphosphatase and phosphoribulokinase (75, 76) can be explored to reduce its oxygen requirement for NADH reoxidation.

The absence of orthologs of the *S. cerevisiae* Aus1 and Pdr11 sterol transporters indicates that, similar to other pre-WGD yeasts (57), *O. parapolymorpha* is most likely unable to import sterols. Due to the incompletely resolved role of cell wall proteins in sterol import in *S. cerevisiae* (77), functional expression of a heterologous system for sterol import may not be a trivial challenge. Alternatively, the importance of sterol uptake by oxygen-limited cultures can be investigated by expression of a heterologous squalene-tetrahymanol cyclase. In *S. cerevisiae* and *K. marxianus*, this modification was recently shown to enable synthesis of the sterol surrogate tetrahymanol and sterol-independent anaerobic growth. Genome-sequence data indicate that pyrimidine synthesis in *O. parapolymorpha* depends on a respiratory-chain-linked dihydroorate dehydrogenase (OpUra9), thus most probably also rendering pyrimidine biosynthesis in this yeast oxygen dependent (78, 79). As previously explored in *Scheffersomyces stipitis* (78), expression of the soluble fumarate-coupled dihydroorotate dehydrogenase from *S. cerevisiae* (ScUra1), may be explored to bypass this oxygen requirement.

We previously showed that, in the thermotolerant yeast *K. marxianus*, an inability to import sterols was the sole feature preventing growth in media supplemented with the know anaerobic growth factors for *S. cerevisiae* (35). The results described in this study on *O. parapolymorpha* underline that oxygen requirements of industrially relevant yeasts, as well as their transcriptional responses to oxygen limitation, show strong quantitative and qualitative differences. In addition to illustrating the need for further fundamental research on oxygen requirements in yeasts and other fungi, this diversity should be considered when selecting host strains for novel anaerobic, yeast-based processes.

## Methods

### Strain maintenance

Yeast strains used in this study were derived from the wild-type *Ogataea parapolymorpha* strain CBS11895 (DL-1) obtained from CBS-KNAW (Westerdijk Fungal Biodiversity Institute, Utrecht, the Netherlands). For propagation and strain maintenance, yeast cultures were grown on yeast peptone dextrose (YPD) media (10 g L^-1^ Bacto yeast extract, 20 g L^-1^ Bacto peptone, 7.5 g L^-1^ glucose) in an Innova shaker incubator (New Brunswick Scientific, Edison, NJ, USA) set at 30 °C and 200 rpm. YP medium was heat sterilized (121 °C for 20 minutes). Stock solutions of concentrated D-glucose were autoclaved separately at 110 °C and added to a concentration of 7.5 g L^-1^. Cryo-stocks were prepared from exponentially growing cultures by addition of glycerol to a final concentration of 30 % (v/v) and aseptically stored at -80 °C.

#### Molecular biology techniques

PCR amplification for cloning was routinely performed with Phusion High Fidelity polymerase (Thermo Fisher Scientific, Waltham, MA, USA) according to the manufacturer’s instructions. DreamTaq polymerase (Thermo Fisher Scientific) was used for diagnostic PCR reactions. Oligonucleotide primers were ordered either desalted or PAGE-purified (Sigma-Aldrich, St. Louis, MO, USA) and are listed in **Table S3**. DNA fragments obtained by PCR amplification were analysed by gel electrophoresis, and when required, purified from agarose gels with the Zymoclean Gel DNA Recovery Kit (Zymo Research, Irvine, CA, USA) according to the manufacturer’s instructions. Prior to purification, template plasmid DNA was removed by FastDigest *DpnI* digestion (Thermo Fisher Scientific). Alternatively, DNA fragments were directly purified from the PCR mix using the GenElute PCR Clean-Up Kit (Sigma-Aldrich). Gibson assemblies were performed with the NEBuilder HiFI DNA Assembly Master mix (New England Biolabs, Ipswich, MA, USA) with the total reaction volume downscaled to 5 µL and a total incubation time of 1 h at 50 °C. *Escherichia coli* XL1-Blue cells were used for transformation, amplification and storage of plasmids. Plasmids were isolated from overnight *E. coli* cultures with the GenElute Plasmid Miniprep kit (Sigma-Aldrich). For diagnostic PCR amplifications, yeast colonies were grown overnight in YPD, after which the genomic DNA was extracted using the LiAc/SDS method (80).

### Plasmid construction

Plasmids used in this study are described in **Table 4**. *O. parapolymorpha* promoters and terminators were chosen based on high transcript levels across a range of specific growth rates (81). Promoter and terminator regions were defined as 800 bp upstream and 300 bp downstream of the coding sequence, respectively. For targeted integration into a genetic locus, 500 bp flanking homology regions were designed to partially delete the target region, without altering promoters (800 bp) or terminators (300 bp) of adjacent genes.

**Table 4.** Plasmids used in this study. Superscripts indicate restriction sites, subscripts indicate gRNA target sequences. Sc: *Saccharomyces cerevisiae*; Op: *Ogataea parapolymorpha*, Ag: *Ashyba gossypii*, Aa: *Arxula adeninivorans*, Sp: *Streptococcus pyogenes*.

| Plasmid | Characteristics | Source |
| --- | --- | --- |
| pUC19 | ori ampR | (85) |
| pUD803 | HR<sub>NDH2-3</sub> AaTEF1p-nat1-ScPHO5t HR<sub>NDH2-3</sub> | (59) |
| pUD1069 | ori amp<sup>R</sup> *gbu1*(HR) OpPMA1p-ScGPP1-OpPMA1t-AgTEF1p-nat1-ScADH1t *gbu1*(HR) | This study |
| pUD1082 | ori ampR *sga1*(HR) OpTEF1p-ScGPD2-OpTEF1t AgTEF1p-hph-AgTEF1t *sga1*(HR) | This study |
| pUDP002 | panARS AgTEF1p-hph-AgTEF1t ScTDH3p<sup>BsaI BsaI</sup>ScCYC1t AaTEF1p-Spcas9-ScPHO5t | (72) |
| pUG6 | ori ampR loxP-kanMX-loxP | (86) |

**Table 4.** Yeast strains used in this study. Sc: *Saccharomyces cerevisiae*; Op: *Ogataea parapolymorpha*

| Genus | Strain | Relevant genotype | Reference |
| --- | --- | --- | --- |
| *S. cerevisiae* | CEN.PK113-7D | MATa URA3 HIS3 LEU2 TRP1 MAL2-8c SUC2 | (87) |
| *O. parapolymorpha* | CBS11895 | - | Westerdijk |
| *O. parapolymorpha* | IMX2119 | *gbu1Δ::OpPMA1p-ScGPP1-OpPMA1t natNT2* | This study |
| *O. parapolymorpha* | IMX2587 | *sga1Δ::OpTEF1p-ScGPD2-OpTEF1t-hph* | This study |
| *O. parapolymorpha* | IMX2588 | *gbu1Δ::OpPMA1p-ScGPP1-OpPMA1t natNT2 sga1Δ::OpTEF1p-ScGPD2-OpTEF1t-hph* | This study |
| *O. parapolymorpha* | IMX2167 | *ndh2-1Δ::kanR ndh2-2Δ::hph ndh2-3::natNT2 gut2Δ::pat* | (59) |

Plasmid pUD1069 (Sc*GPP1*) was constructed by Gibson assembly from fragments with 20 bp terminal sequence overlaps. The *GPP1* coding sequence was PCR amplified from genomic DNA of *S. cerevisiae* CEN.PK113-7D with primers 15183 and 15814 thereby introducing 20 bp overlaps. The promoter and terminator of the endogenous *O. parapolymorpha PMA1* were amplified from genomic DNA using primers 15185/15186 and, 15191/15193 respectively. The natNT2 marker (Ag*TEF1*p-nat1-Sc*ADH1*t) from *Streptomyces noursei* (82–84) was PCR amplified from pUD803 (59) with primers 15187 and 15188. Up- and downstream 500 bps homology flanks to the *gbu1* locus were PCR amplified from genomic DNA of *O. parapolymorpha* using primers 15192/15196 and 15195/15198 respectively. The pUG6 backbone was PCR amplified using primers 15189 and 15190. Fragments were mixed in equimolar amounts and Gibson assembled resulting in plasmid pUD1069. Correct plasmid assembly was verified by restriction digestion and diagnostic PCR amplification with primers 15224, 15225, 15226, 15227, 15228, 15229, 15230, 15231 and 15232.

For heterologous expression of Sc*GPD2*, its coding sequence was PCR amplified from genomic DNA of CEN.PK113-7D with primers 15745 and 15744, resulting in the addition of 20 bp homology flanks for Gibson assembly. The Op*TEF1* promoter and Op*TEF1* terminator were PCR amplified from genomic DNA of *O. parapolymorpha* CBS11895 with primers 15749/15748 and 15746/15747, respectively. Upstream and downstream 800-bp recombination flanks for integration of the *SGA1* integration locus were PCR amplified with primers 15740/15739 and 15735/15736, respectively. A *Klebsiella pneumoniae hph* (Hyg^R^) marker cassette, including the Ag*TEF1* promoter and Ag*TEF1* terminator from *Ashbya gossypii*, was PCR amplified from pUDP002 (72) with primers 1312 and 15743. The pUC19 backbone was PCR amplified with primers 15738 and 15737. The resulting 6 fragments were mixed in in equimolar amounts and Gibson assembled. The resulting plasmid pUD1082 (Sc*GPD2*) was verified by restriction digestion.

### Strain construction

*O. parapolymorpha* strains were transformed by electroporation of freshly prepared electrocompetent cells (72). Yeast cells were selected after transformation on YPD agar containing hygromycin B (300 µg mL^-1^) or nourseothricin (100 µg mL^-1^). Strains IMX2119, IMX2587 and IMX2588 were constructed using the split-marker integration approach (88), with an approximately 480 bp overlapping homology sequences for marker recombination and genome integration. The natNT2 split-marker fragments for integration of a Sc*GPP1* expression cassette into the Op*GBU1* locus were constructed by PCR amplification with primers 15192, 15194, 15196 and 15197 from pUD1069 (Sc*GPP1*) resulting in two integration fragments with a homologous sequence overlap. Similarly, the hph split-marker fragments for integration of the Sc*GPD2* cassette were constructed by PCR amplification of a pUD1082 (Sc*GPD2*) fragment with primers 15740, 15741, 15742 and 15736. The chromosomal integration of the split-marker fragment was verified by diagnostic PCR. Diagnostic PCR of the Op*GBU1* integration locus was performed with primers 15192, 15197, 15233 and 15234, and integration in the Op*SGA1* integration locus was verified with primers 15894, 15748, 15895.

### Bioreactor cultivation

Aerobic batch and chemostat cultures for physiological characterization of *O. parapolymorpha* strains CBS11895 and IMX2167 strains were grown in 2 L bioreactors (Applikon Biotechnology, Delft, the Netherlands) with 1.0 L working volume. Cultures were grown on synthetic medium with vitamins (89), supplemented with glucose and with sterile antifoam pluronic acid 6100 PE (0.2 g L^-1^, BASF, Ludwigshafen, Germany). Mineral salt solutions were sterilized by autoclaving at 121 °C for 20 min. Glucose solutions were prepared and autoclaved separately at 110 °C for 20 min and added to a final concentration of 7.5 g L^-1^.

Inocula for bioreactor cultures were prepared by harvesting an exponential growing shake-flask culture by centrifugation and washing once with sterile demineralized water. Cultures were continuously stirred at 800 rpm, maintained at 30 °C and sparged with air (O_2_ 21·10^4^ ppm) at a volumetric rate of 0.5 L min^-1^ (0.5 vvm) to maintain a high dissolved oxygen concentration. After CO_2_ emission in the batch cultivation phase had reached a maximum and started to decline sharply, a constant medium feed and continuous effluent removal were initiated to maintain a constant working volume.

Oxygen-limited chemostat cultures of *O. parapolymorpha* strains were grown in 2 L bioreactors (Applikon) with 1.2 L working volume on synthetic medium with vitamins and urea as nitrogen source (90). A stock solution of urea was filter sterilized (0.2 µm) and added to the sterile media. Before autoclaving, bioreactors were checked for gas leakage by submersion in water while applying a 0.3 bar overpressure. Chemostats were started as aerobic batch cultures to obtain a sufficiently high initial cell density. The glucose concentration in the aerobic batch phase was 1.5 g L^-1^ while the chemostat feed media contained 20 g L^-1^ glucose for the oxygen-limited and 7.5 g L^-1^ for the aerobic reference cultures.

The anaerobic growth factors ergosterol (10 mg L^-1^, Sigma Aldrich), Tween 80 (84 mg L^-1^, Sigma Aldrich) were solubilized in pure ethanol (671 µg L^-1^, Sigma Aldrich). Sterile anaerobic growth factors ergosterol and Tween 80 were prepared as described previously (39), to prevent excessive foam formation while still supporting maximal growth the Tween 80 concentration was reduced 5 fold compared to anaerobic *S. cerevisiae* reference cultures. For aerobic chemostat cultures, Tween 80 was omitted to prevent excessive foam formation and the glucose concentration was adjusted to 7.5 g L^-1^. Dissolved gas concentrations in the bioreactor cultures were maintained by sparging with either air (21·10^4^ ppm O_2_) or a gas mixture of N_2_/Air (840 ppm O_2_) at a volumetric rate of 0.5 L min^-1^ (0.4 vvm). Bioreactors were equipped with Fluran tubing and viton O-rings to minimize oxygen diffusion. The glass medium reservoir was equipped with Norprene tubing and continuously sparged with pure nitrogen gas.

### Analytical methods

Off-gas analysis, biomass dry weight measurements, optical density measurements, metabolite HPLC analysis of culture supernatants and correction for ethanol evaporation in bioreactor experiments were performed as described previously (35). Rates of substrate consumption and metabolite production were determined from glucose and metabolite concentrations in steady-state cultures, analyzed after rapid quenching of culture samples (91). Carbon and degree of reduction balances were calculated based on concentrations of relevant components in medium feed, culture samples and in- and out-going gas streams. Calculations were based on an estimated degree of reduction of biomass as described in (92).

### RNA extraction, isolation, sequencing and transcriptome analysis

Biomass samples from bioreactor batch and chemostat cultures were directly sampled into liquid nitrogen to prevent mRNA turnover (93) and stored at -80 °C before extraction and isolation. Batch cultures were sampled in mid-exponential phase when approximately 75 % of the initial glucose concentration was still unused (81). Processing of RNA for long-term storage and isolation was performed as described previously (35). The quality of isolated RNA was analyzed with an Agilent Tapestation (Agilent Technologies, CA, USA) using RNA screen tape (Agilent). RNA concentrations were measured with a Qubit RNA BR assay kit (Thermo Fischer Scientific). The TruSeq Stranded mRNA LT protocol (Illumina, San Diego, CA, USA) was used to generate RNA libraries for paired-end sequencing by Macrogen (Macrogen Europe, Amsterdam, the Netherlands) with a read-length of 151 bp on a NovaSeq sequencer (Illumina).

To refine the genome annotation of *O. parapolymorpha* CBS11895, pooled RNAseq libraries were first used to perform *de novo* transcriptome assembly using Trinity (v2.8.3)(94) and the used for the PASA pipeline (95) as implemented in funannotate (96). RNA reads from *O. parapolymorpha* wild-type batch and chemostat (81) were downloaded from NCBI (www.ncbi.nlm.nih.gov) with the gene expression omnibus accession GSE140480.

RNA reads were mapped to the genome of strain CBS11895 (81) using bowtie (v1.2.1.1) (97) and alignments were filtered and sorted using samtools (v1.3.1)(98) as described previously (35). Reads were counted with featureCounts (v1.6.0)(99) of which both pairs of the paired-end reads were aligned to the same chromosome. edgeR (v3.28.1)(100) was used to perform differential gene expression and genes with fewer than 10 reads per million in all conditions were eliminated from subsequent analysis. Counts were normalized using the trimmed mean of M values (TMM)(101) method and the dispersion was estimated using generalized linear models. Differential expression was calculated using a log ratio test adjusted with the Benjamini-Hochberg method. Absolute log 2 fold-change values (>2), false discovery rate (< 0.5) and *p*-value (<0.05) were used a significance cutoffs.

Gene set analysis (GSA) based on gene ontology (GO) terms with Piano (v2.4.0)(58) was used for functional interpretation of differential gene expression profiles. Interproscan (102) was used to assign GO terms to the genome annotation of *O. parapolymorpha*. Co-ortholog groups of genes were generated with ProteinOrtho5 (103) as implemented in the funannotate pipeline and used to homogenize GO terms for co-ortholog groups as described previously (35). Gene set analysis was performed with Piano (v2.4.0)(58) and gene statistics were calculated with Stouffer, Wilcoxon rank-sum test and reporter methods as implemented in Piano. Consensus gene level statistics were obtained by *p*-value and rank aggregation and considered significant based on absolute log 2 fold-change values > 1. ComplexHeatmap (v2.4.3)(104) was used to visualize the differentially expressed genes, including those shared with a previous study (55).

To interpret the GO-term based gene set analysis between three yeast species in response to oxygen limitation, hierarchical clustering (complete method and Euclidian distance) in R (105) was performed on GO-terms from biological process category. Clustering was based on the number of overlapping distinct directionality *p*-values in the three yeast species with a significance *p*-value cut-off of 0.01.

### Sequence homology search

*S. cerevisiae* protein sequences were used as query for searching whole genome sequences of 16 *Ogataea* species, *K. marxianus, Candida arabinofermentans* and *B. bruxellensis* with tblastn (blast.ncbi.nlm.nih.gov)(106). Significance was based on alignment criteria with an e-value (< 10^−7^), alignment coverage (>70%) and nucleotide percent identity of (>50%). Blast results were mapped to a subtree of selected yeast species in the fungal phylum Ascomycota (48) using Treehouse (107) to subset the phylogenetic tree.

## Data availability

Data presented in all figures in this work are available at the www.data.4TU.nl repository doi: 10.4121/14270138.

Raw sequencing data that supports this study are available from the NCBI website (www.ncbi.nlm.nih.gov/geo/) with the BioProject PRJNA717220.

## Code availability

The codes that were used to generate the results obtained in this study are archived in a Gitlab repository (https://gitlab.tudelft.nl/rortizmerino/opar_anaerobic).

## Author Contributions

WD, and JTP designed the study and wrote the manuscript. WD performed molecular cloning with the support of MG and RB. WD, HJ and CM performed the bioreactor experiments with the support of AK. RO contributed the transcriptome analysis and genome annotation.

## Acknowledgements

We thank Hans van Dijken, Mark Bisschops and Jonna Bouwknegt for fruitful discussions. We thank Erik de Hulster for fermentation support and Nikolai Gyurchev and Janine Nijenhuis for their input.

## Supplementary Material

**Supplementary Figure 1.**
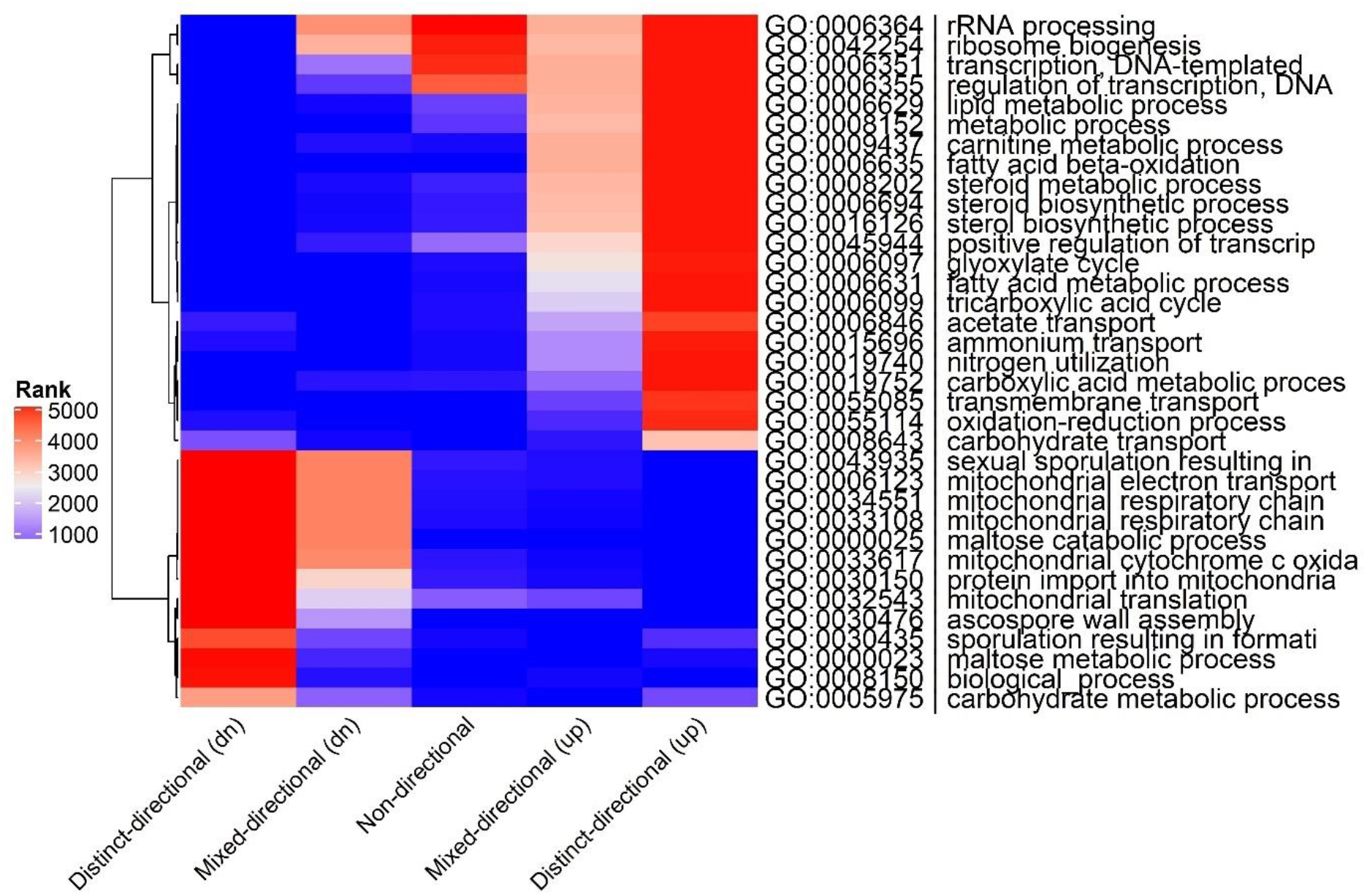
Consensus biological process GO term enrichment for *S. cerevisiae* contrast 21. GO terms are clustered according to their rank. See legend of Fig. 3 for experimental details.

**Supplementary Figure 2.**
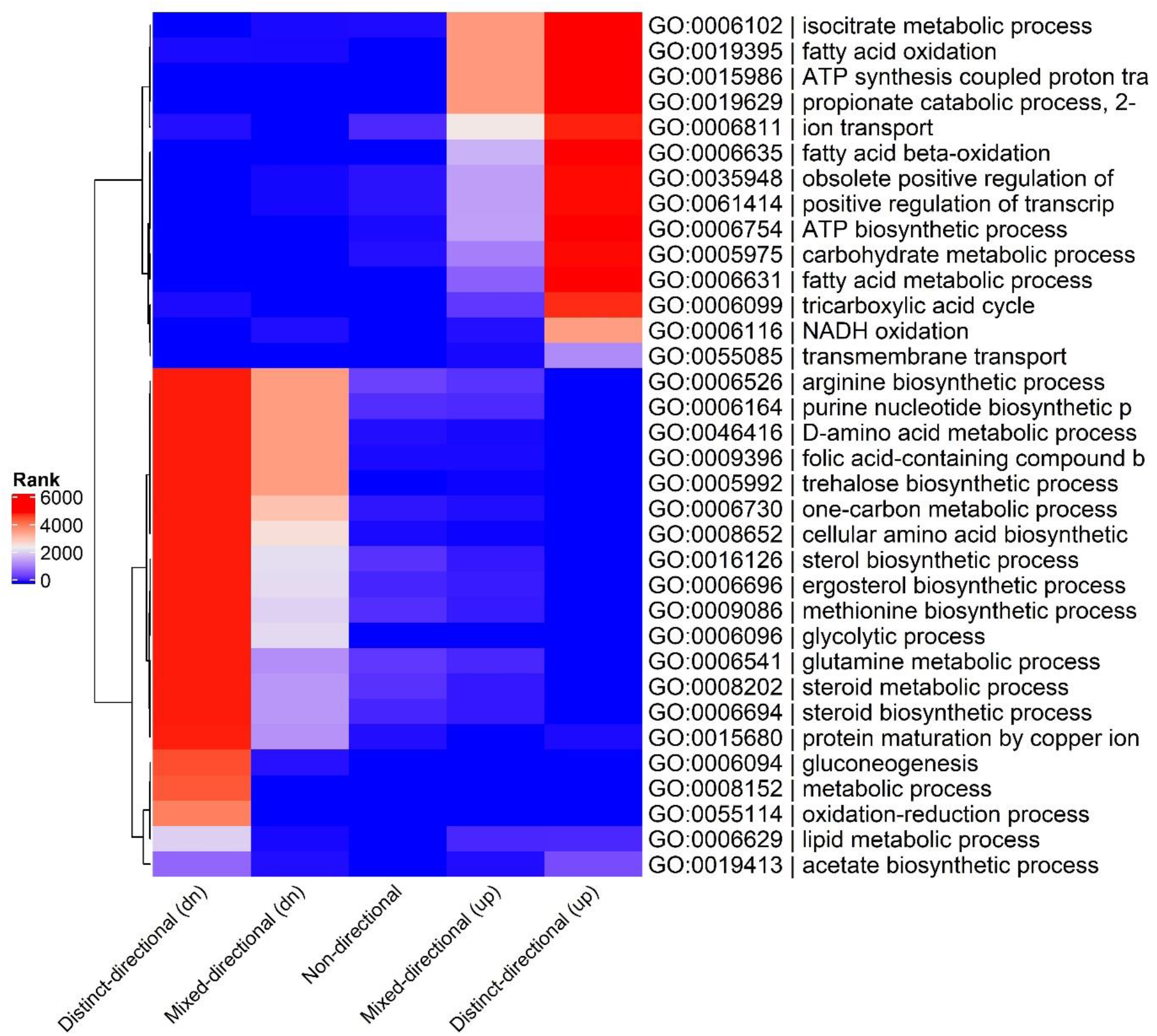
Consensus biological process GO term enrichment for *K. marxianus* contrast 21. GO terms are clustered according to their rank. See legend of **Fig. 3** for experimental details.

**Supplementary Figure 3.**
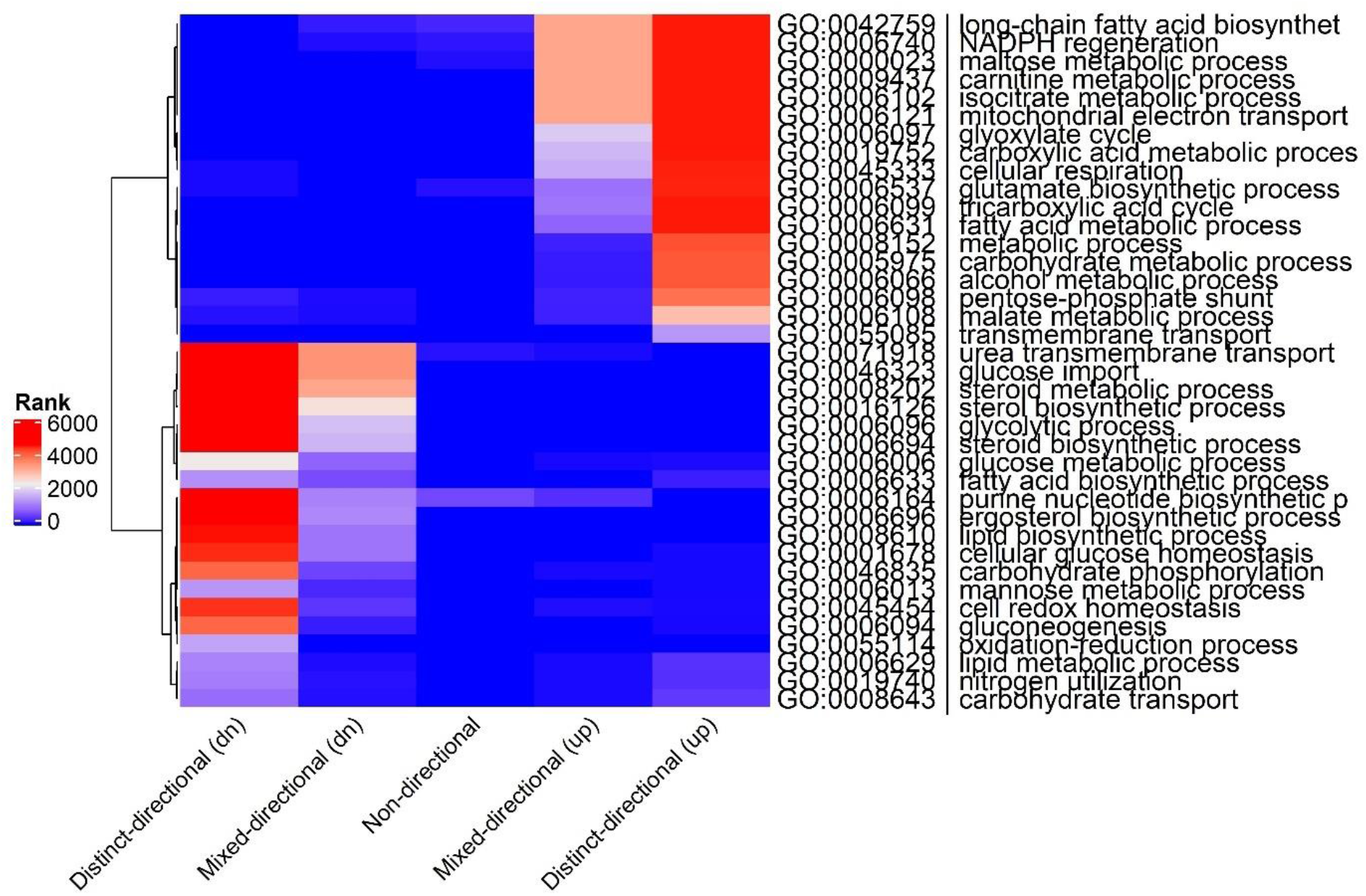
Consensus biological process GO term enrichment for *O. parapolymorpha* contrast 21. GO terms are clustered according to their rank. See legend of **Fig. 3** for experimental details.

**Supplementary Figure 4.**
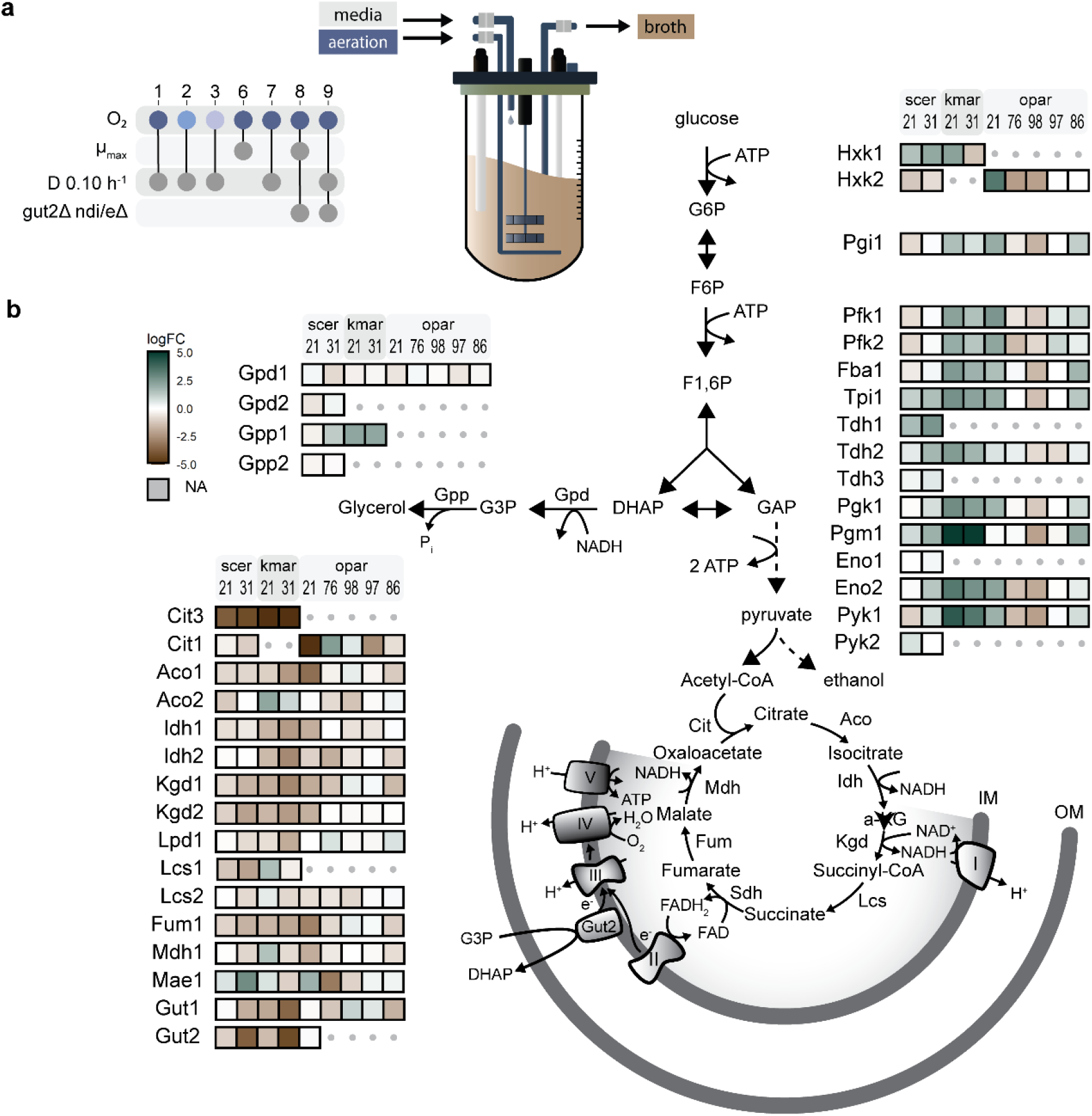
Different transcriptional regulation of NADH reduction in *O. parapolymorpha, K. marxianus, S. cerevisiae*. (**A**) For each aeration regime (1 to 3) RNAseq was performed of chemostat cultures (dilution rate 0.10 h^-1^) for *O. parapolymorpha* CBS11895 and, *K. marxianus* CBS6556 and *S. cerevisiae* CEN.PK113-7D where possible (contrast 31 was not possible for *O. parapolymorpha*). RNAseq data of *S. cerevisiae* and *K. marxianus* was reanalyzed from (35) which included anaerobic-growth factor supplementation regimes (4 to 5). RNAseq was performed on independent replicate batch and chemostat cultures of *O. parapolymorpha* CBS11895 and IMX2167 (*gut2*Δ *ndi/e*Δ) in aerobic conditions (condition/regime 6 to 9). (**B**) Biochemical reactions are represented by arrows and multiple reactions by dashed arrows. Boxes with colors indicate up-(blue-green) or downregulation (brown) with color intensity indicating the log 2 fold change (logFC) with color range capped to a maximum value of 5. Reactions are annotated with the corresponding *S. cerevisiae* ortholog name and grey dots indicate lack of an ortholog in the respective yeast. Abbreviations used; pentose-phosphate pathway (PPP), glucose-6-phosphate (G6P), fructose-6-phosphate (F6P), fructose-1,6-bisphosphate (F1,6P), dihydroxyacetone phosphate (DHAP), glycerol-3P (G3P), glyceraldehyde phosphate (GAP), inner-mitochondrial membrane (IM), OM outer-mitochondrial membrane, respiratory chain complex’s annotated with corresponding roman numerals.

**Supplementary Figure 5.**
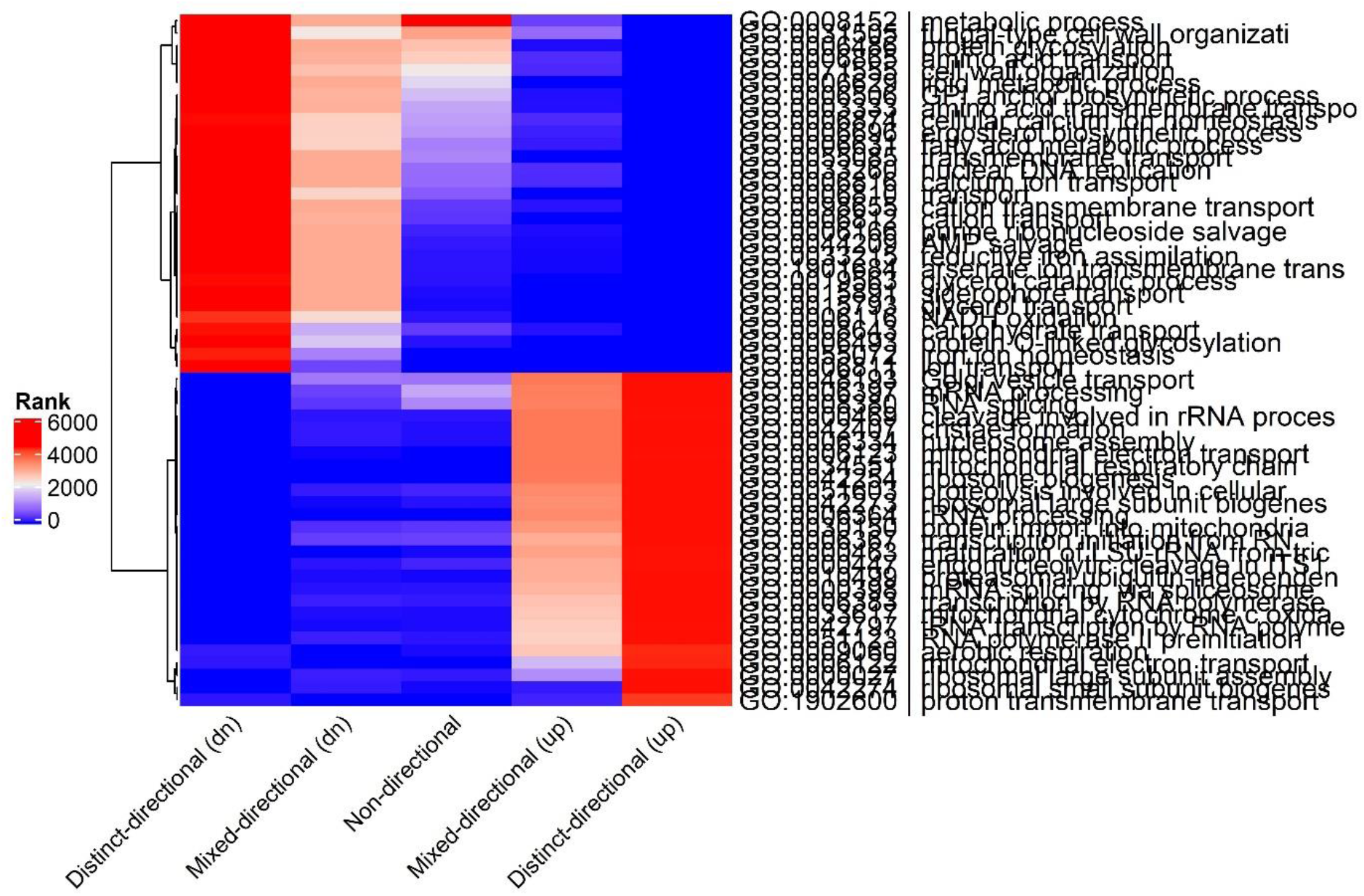
Consensus biological process GO term enrichment for *O. parapolymorpha* contrast 61. GO terms are clustered according to their rank. See legend of **Fig. S4** for experimental details.

**Supplementary Table 1.** Transcriptional response of phosphatases in *O. parapolymorpha* wild-type to *gut2*Δ *nde/i1-3*Δ knockout in aerobic batch cultures. Log fold-change (logFC), log counts per million (logCPM), gene identifiers used in the genome annotations and *S. cerevisiae* S288C orthologs and three-letter abbreviation (standard name).

| logFC | logCPM | PValue | FDR | Geneid | DL.1 | S288C | Standard Name |
| --- | --- | --- | --- | --- | --- | --- | --- |
| -1.61 | 5.74 | 2.06E-19 | 1.11E-18 | OPAR_004591 | HPODL_02465 | YGR208W | SER2 |
| -1.56 | 3.52 | 1.10E-08 | 2.56E-08 | OPAR_004515 | HPODL_01991 |  |  |
| 1.39 | 7.90 | 1.51E-23 | 1.07E-22 | OPAR_002912 | HPODL_03896 | YKL212W | SAC1 |
| -1.37 | 6.68 | 2.73E-18 | 1.36E-17 | OPAR_005066 | HPODL_02910 | YOR283W |  |
| 1.36 | 5.25 | 9.20E-12 | 2.80E-11 | OPAR_002432 | HPODL_03461 | YJR110W | YMR1 |
| 1.29 | 6.58 | 6.85E-21 | 4.09E-20 | OPAR_000036 | HPODL_00031 | YFR028C | OAF3 |
| 1.19 | 5.46 | 2.48E-06 | 4.74E-06 | OPAR_004387 | HPODL_01871 | YDL230W | PTP1 |
| -0.99 | 7.74 | 1.28E-12 | 4.11E-12 | OPAR_003314 | HPODL_04287 | YDL236W | PHO13 |
| 0.93 | 5.77 | 2.55E-09 | 6.27E-09 | OPAR_001025 | HPODL_04916 | YBR204C | LDH1 |
| 0.73 | 6.09 | 5.08E-06 | 9.40E-06 | OPAR_002585 | HPODL_03597 | YBR276C | PPS1 |
| 0.64 | 5.10 | 0.011907 | 0.01586 | OPAR_000879 | HPODL_04782 | YDL057W |  |
| -0.64 | 6.01 | 0.000114 | 0.000186 | OPAR_005045 | HPODL_02888 | YNL128W | TEP1 |
| 0.63 | 5.72 | 0.000104 | 0.00017 | OPAR_001327 | HPODL_00279 | YDR481C | PHO8 |
| 0.58 | 7.20 | 1.47E-05 | 2.60E-05 | OPAR_001974 | HPODL_00894 | YBR092C | PHO3 |
| 0.52 | 7.69 | 8.32E-05 | 0.000139 | OPAR_003977 | HPODL_01484 | YLR019W | PSR2 |
| 0.48 | 6.67 | 0.001628 | 0.002382 | OPAR_002901 | HPODL_03887 | YOR155C | ISN1 |
| 0.45 | 5.71 | 0.003449 | 0.004877 | OPAR_004178 | HPODL_01675 | YNL217W | PPN2 |
| -0.43 | 8.41 | 0.00185 | 0.002689 | OPAR_005130 | HPODL_02972 | YNL010W | PYP1 |
| 0.38 | 6.44 | 0.004739 | 0.006577 | OPAR_001369 | HPODL_00320 |  |  |
| -0.28 | 4.14 | 0.256548 | 0.289337 | OPAR_002548 | HPODL_03567 | YMR087W | PDL32 |
| -0.28 | 6.41 | 0.066792 | 0.081355 | OPAR_001685 | HPODL_00615 | YHR004C | NEM1 |
| 0.18 | 6.02 | 0.220151 | 0.25152 | OPAR_000296 | HPODL_02098 |  |  |
| 0.12 | 7.52 | 0.372061 | 0.408252 | OPAR_002280 | HPODL_03321 | YLR377C | FBP1 |
| 0.12 | 6.55 | 0.366906 | 0.402931 | OPAR_002299 | HPODL_03339 | YNL108C |  |

**Supplementary Table 2.**
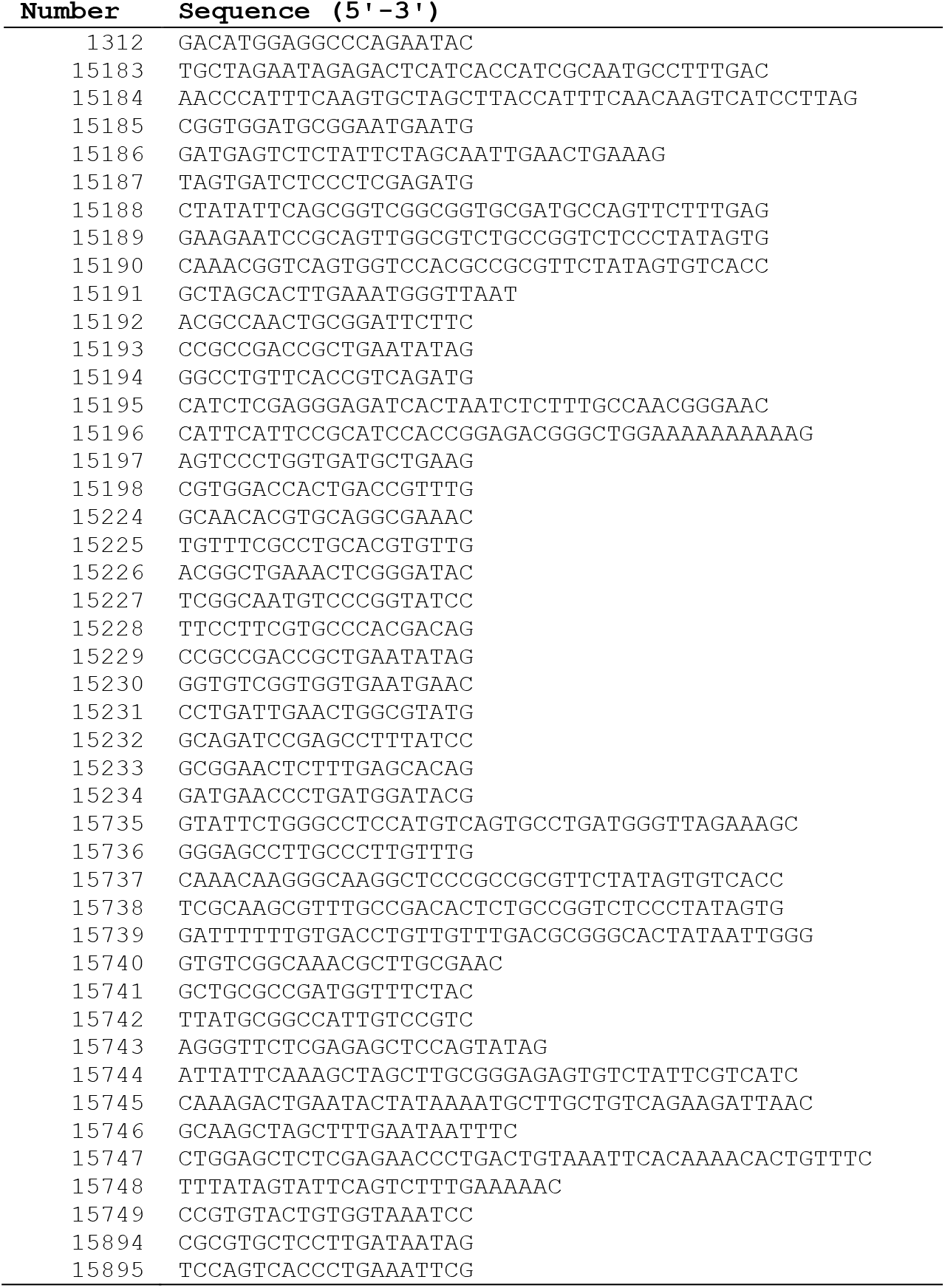
Primers used in this study.

## References

1. Weusthuis RA, Lamot I, van der Oost J, Sanders JPM. 2011. Microbial production of bulk chemicals: Development of anaerobic processes. Trends Biotechnol 29:153–158.

2. Thorwall S, Schwartz C, Chartron JW, Wheeldon I. 2020. Stress-tolerant non-conventional microbes enable next-generation chemical biosynthesis. Nat Chem Biol 16:113–121.

3. 2020. Annual World Fuel Ethanol Production. Renew Fuels Assoc.

4. Jansen MLA, Bracher JM, Papapetridis I, Verhoeven MD, de Bruijn H, de Waal PP, van Maris AJA, Klaassen P, Pronk JT. 2017. Saccharomyces cerevisiae strains for second-generation ethanol production: from academic exploration to industrial implementation. FEMS Yeast Res 17:1–20.

5. Favaro L, Jansen T, van Zyl WH. 2019. Exploring industrial and natural Saccharomyces cerevisiae strains for the bio-based economy from biomass: the case of bioethanol. Crit Rev Biotechnol 39:800–816.

6. Basso LC, Basso TO, Rocha SN. 2011. Ethanol production in Brazil: the industrial process and its impact on yeast fermentation. Biofuel Prod Dev Prospect 1530:85–100.

7. van Maris AJA, Abbott DA, Bellissimi E, van den Brink J, Kuyper M, Luttik MAH, Wisselink HW, Scheffers WA, van Dijken JP, Pronk JT. 2006. Alcoholic fermentation of carbon sources in biomass hydrolysates by Saccharomyces cerevisiae: Current status. Antonie van Leeuwenhoek, Int J Gen Mol Microbiol 90:391–418.

8. Thomas KC, Ingledew WM. 1992. Production of 21% (v/v) ethanol by fermentation of very high gravity (VHG) wheat mashes. J Ind Microbiol 10:61–68.

9. Della-Bianca BE, Basso TO, Stambuk BU, Basso LC, Gombert AK. 2013. What do we know about the yeast strains from the Brazilian fuel ethanol industry? Appl Microbiol Biotechnol 97:979–991.

10. Lopes ML, Paulillo SC de L, Godoy A, Cherubin RA, Lorenzi MS, Giometti FHC, Bernardino CD, Amorim Neto HB de, Amorim HV de. 2016. Ethanol production in Brazil: a bridge between science and industry. Brazilian J Microbiol 47:64–76.

11. Laman Trip DS, Youk H. 2020. Yeasts collectively extend the limits of habitable temperatures by secreting glutathione. Nat Microbiol 5:943–954.

12. Althuri A, Chintagunta AD, Sherpa KC, Banerjee R. 2018. Simultaneous Saccharification and Fermentation of Lignocellulosic Biomass, p. 265–285.In Kumar, S, Sani, RK (eds.), Biorefining of Biomass to Biofuels: Opportunities and Perception. Springer International Publishing, Cham.

13. Costa DA, De Souza CJA, Costa PS, Rodrigues MQRB, Dos Santos AF, Lopes MR, Genier HLA, Silveira WB, Fietto LG. 2014. Physiological characterization of thermotolerant yeast for cellulosic ethanol production. Appl Microbiol Biotechnol 98:3829–3840.

14. Alvira P, Tomás-Pejó E, Ballesteros M, Negro MJ. 2010. Pretreatment technologies for an efficient bioethanol production process based on enzymatic hydrolysis: A review. Bioresour Technol 101:4851–4861.

15. Shirkavand E, Baroutian S, Gapes DJ, Young BR. 2016. Combination of fungal and physicochemical processes for lignocellulosic biomass pretreatment. Renew Sustain Energy Rev 54:217–234.

16. Cruz LAB, Hebly M, Duong G-H, Wahl SA, Pronk JT, Heijnen JJ, Daran-Lapujade P, van Gulik WM. 2012. Similar temperature dependencies of glycolytic enzymes: an evolutionary adaptation to temperature dynamics? BMC Syst Biol 6:151.

17. Li G, Hu Y, Jan Zrimec, Luo H, Wang H, Zelezniak A, Ji B, Nielsen J. 2021. Bayesian genome scale modelling identifies thermal determinants of yeast metabolism. Nat Commun 12:190.

18. Caspeta L, Nielsen J. 2015. Thermotolerant Yeast Strains Adapted by Laboratory Evolution Show Trade-Off at Ancestral Temperatures and Preadaptation to Other. MBio 6:1–9.

19. Caspeta L, Chen Y, Ghiaci P, Feizi A, Buskov S, Hallström BM, Petranovic D, Nielsen J. 2014. Altered sterol composition renders yeast thermotolerant. Science (80-) 346:75–78.

20. Kurtzman CP. 2011. A new methanol assimilating yeast, Ogataea parapolymorpha, the ascosporic state of Candida parapolymorpha. Antonie van Leeuwenhoek, Int J Gen Mol Microbiol 100:455–462.

21. Kurylenko OO, Ruchala J, Hryniv OB, Abbas CA, Dmytruk K V., Sibirny AA. 2014. Metabolic engineering and classical selection of the methylotrophic thermotolerant yeast Hansenula polymorpha for improvement of high-temperature xylose alcoholic fermentation. Microb Cell Fact 13:1–10.

22. Hong J, Wang Y, Kumagai H, Tamaki H. 2007. Construction of thermotolerant yeast expressing thermostable cellulase genes. J Biotechnol 130:114–123.

23. Visser W, Scheffers WA, Batenburg-Van der Vegte WH, Van Dijken JP. 1990. Oxygen requirements of yeasts. Appl Environ Microbiol 56:3785–3792.

24. Merico A, Sulo P, Piškur J, Compagno C. 2007. Fermentative lifestyle in yeasts belonging to the Saccharomyces complex. FEBS J 274:976–989.

25. Merico A, Galafassi S, Piškur J, Compagno C. 2009. The oxygen level determines the fermentation pattern in Kluyveromyces lactis. FEMS Yeast Res 9:749–756.

26. Wolfe KH, Shields DC. 1997. Molecular evidence for an ancient duplication of the entire yeast genome. Nature 387:708–713.

27. Blomqvist J, Nogue VS, Gorwa-Grauslund M, Passoth V. 2012. Physiological requirements for growth and competitveness of Dekkera bruxellensis under oxygen limited or anaerobic conditions. Yeast 29:265–274.

28. Snoek ISI, Steensma HY. 2006. Why does Kluyveromyces lactis not grow under anaerobic conditions? Comparison of essential anaerobic genes of Saccharomyces cerevisiae with the Kluyveromyces lactis genome. FEMS Yeast Res 6:393–403.

29. Stukey JE, McDonough VM, Martin CE. 1989. Isolation and Characterization of OLE1, a Gene Affecting Fatty Acid Desaturation from Saccharomyces cerevisiae. J Biol Chem 264:16537– 16544.

30. Henneberry AL, Sturley SL. 2005. Sterol homeostasis in the budding yeast, Saccharomyces cerevisiae. Semin Cell Dev Biol 16:155–161.

31. Andreasen AA, Stier TJB. 1953. Anaerobic nutrition of Saccharomyces cerevisiae II. Unsaturated fatty acid requirement for growth in a defined medium. J Cell Physiol 43:271– 281.

32. Langkjær RB, Cliften PF, Johnston M, Piškur J. 2003. Yeast genome duplication was followed by asynchronous differentiation of duplicated genes. Nature 421:848–852.

33. Nagy M, Lacroute F, Thomas D. 1992. Divergent evolution of pyrimidine biosynthesis between anaerobic and aerobic yeasts. Proc Natl Acad Sci U S A 89:8966–8970.

34. Riley R, Haridas S, Wolfe KH, Lopes MR, Hittinger CT, Göker M, Salamov AA, Wisecaver JH, Long TM, Calvey CH, Aerts AL, Barry KW, Choi C, Clum A, Coughlan AY, Deshpande S, Douglass AP, Hanson SJ, Klenk H-P, LaButti KM, Lapidus A, Lindquist EA, Lipzen AM, Meier-Kolthoff JP, Ohm RA, Otillar RP, Pangilinan JL, Peng Y, Rokas A, Rosa CA, Scheuner C, Sibirny AA, Slot JC, Stielow JB, Sun H, Kurtzman CP, Blackwell M, Grigoriev I V, Jeffries TW. 2016. Comparative genomics of biotechnologically important yeasts. Proc Natl Acad Sci U S A 113:9882–9887.

35. Dekker WJC, Ortiz-Merino RA, Kaljouw A, Battjes J, Wiering FW, Mooiman C, de la Torre P. 2021. Engineering the thermotolerant industrial yeast Kluyveromyces marxianus for anaerobic growth. bioRxiv 1–16.

36. Suh SO, Zhou JJ. 2010. Methylotrophic yeasts near Ogataea (Hansenula) polymorpha: A proposal of Ogataea angusta comb. nov. and Candida parapolymorpha sp. nov. FEMS Yeast Res 10:631–638.

37. Verduyn C, Stouthamer AH, Scheffers WA, van Dijken JP. 1991. A theoretical evaluation of growth yields of yeasts. Antonie Van Leeuwenhoek 59:49–63.

38. Andreasen AA, Stier TJB. 1953. Anaerobic nutrition of Saccharomyces cerevisiae I. Ergosterol requirement for growth in a defined medium. J Cell Physiol 41:23–26.

39. Dekker WJC, Wiersma SJ, Bouwknegt J, Mooiman C, Pronk JT. 2019. Anaerobic growth of Saccharomyces cerevisiae CEN.PK113-7D does not depend on synthesis or supplementation of unsaturated fatty acids. FEMS Yeast Res 19.

40. Scheffers WA. 1966. Stimulation of Fermentation in Yeasts by Acetoin and Oxygen. Nature 210:533–536.

41. Weusthuis RA, Visser W, Pronk JT, Scheffers WA, Van Dijken JP. 1994. Effects of oxygen limitation on sugar metabolism in yeasts: A continuous-culture study of the Kluyver effect. Microbiology 140:703–715.

42. van Dijken JP, Scheffers WA. 1986. Redox balances in the metabolism of sugars by yeasts. FEMS Microbiol Lett 32:199–224.

43. Bakker BM, Overkamp KM, Van Maris AJA, Kötter P, Luttik MAH, Van Dijken JP, Pronk JT. 2001. Stoichiometry and compartmentation of NADH metabolism in Saccharomyces cerevisiae. FEMS Microbiol Rev 25:15–37.

44. Scheffers WA. 1963. Effects of oxygen and acetoin on fermentation in yeasts. Antonie Van Leeuwenhoek 29:325–326.

45. Norbeck J, Påhlman A, Akhtar N, Blomberg A, Adler L. 1996. Purification and Characterization of Two Isoenzymes of DL -Glycerol-3-phosphatase from Saccharomyces cerevisiae. J Biol Chem 271:13875–13881.

46. Ansell R, Granath K, Hohmann S, Thevelein JM, Adler L. 1997. The two isoenzymes for yeast NAD+-dependent glycerol 3-phosphate dehyrogenase encoded by GPD1 and GPD2 have distinct roles in osmoadaptation and redox regulation. EMBO J 16:2179–2187.

47. Albertyn J, Tonder A van, Prior BA. 1992. Purification and characterization of glycerol-3-phosphate dehydrogenase of Saccharomyces cerevisiae. FEBS J 308:130–132.

48. Shen X-X, Steenwyk JL, LaBella AL, Opulente DA, Zhou X, Kominek J, Li Y, Groenewald M, Hittinger CT, Rokas A. 2020. Genome-scale phylogeny and contrasting modes of genome evolution in the fungal phylum Ascomycota. Sci Adv 6:eabd0079.

49. Wijsman MR, van Dijken JP, van Kleeff BHA, Scheffers WA. 1984. Inhibition of fermentation and growth in batch cultures of the yeast Brettanomyces intermedius upon a shift from aerobic to anaerobic conditions (Custers effect). Antonie Van Leeuwenhoek 50:183–192.

50. Galafassi S, Capusoni C, Moktaduzzaman M, Compagno C. 2013. Utilization of nitrate abolishes the “Custers effect” in Dekkera bruxellensis and determines a different pattern of fermentation products. J Ind Microbiol Biotechnol 40:297–303.

51. Athenstaedt K, Weys S, Paltauf F, Daum G. 1999. Redundant systems of phosphatidic acid biosynthesis via acylation of glycerol-3-phosphate or dihydroxyacetone phosphate in the yeast Saccharomyces cerevisiae. J Bacteriol 181:1458–1463.

52. Larsson C, Pahlman I, Ansell R, Rigoulet M, Adler L, Gustafsson L. 1998. The importance of the glycerol 3-phosphate shuttle during aerobic growth of Saccharomyces cerevisiae. Yeast 14:347–357.

53. Rigoulet M, Aguilaniu H, Avéret N, Bunoust O, Camougrand N, Grandier-Vazeille X, Larsson C, Pahlman I-L, Manon S, Gustafsson L. 2004. Organization and regulation of the cytosolic NADH metabolism in the yeast Saccharomyces cerevisiae. Mol Cell Biochem 256:73–81.

54. Overkamp KM, Bakker BM, Kötter P, van Tuijl A, de Vries S, van Dijken JP, Pronk JT. 2000. In Vivo Analysis of the Mechanisms for Oxidation of Cytosolic NADH by Saccharomyces cerevisiae Mitochondria. J Bacteriol 182:2823–2830.

55. Tai SL, Boer VM, Daran-Lapujade P, Walsh MC, De Winde JH, Daran JM, Pronk JT, Tai SL, Boer VM, Daran-Lapujade P, Walsh MC, De Winde JH, Daran JM, Pronk JT. 2005. Two-dimensional transcriptome analysis in chemostat cultures: Combinatorial effects of oxygen availability and macronutrient limitation in Saccharomyces cerevisiae. J Biol Chem 280:437–447.

56. Meijer MMC, Boonstra J, Verkleij AJ, Verrips CT. 1998. Glucose repression in Saccharomyces cerevisiae is related to the glucose concentration rather than the glucose flux. J Biol Chem 273:24102–24107.

57. Seret ML, Diffels JF, Goffeau A, Baret P V. 2009. Combined phylogeny and neighborhood analysis of the evolution of the ABC transporters conferring multiple drug resistance in hemiascomycete yeasts. BMC Genomics 10:459.

58. Väremo L, Nielsen J, Nookaew I. 2013. Enriching the gene set analysis of genome-wide data by incorporating directionality of gene expression and combining statistical hypotheses and methods. Nucleic Acids Res 41:4378–4391.

59. Juergens H, Mielgo-Gómez Á, Godoy-Hernández A, ter Horst J, Nijenhuis JM, McMillan DGG, Mans R. 2021. Physiological relevance, localization and substrate specificity of the alternative (type II) mitochondrial NADH dehydrogenases of Ogataea parapolymorpha. bioRxiv 2021.04.28.441406.

60. Geertman JMA, van Maris AJA, van Dijken JP, Pronk JT. 2006. Physiological and genetic engineering of cytosolic redox metabolism in Saccharomyces cerevisiae for improved glycerol production. Metab Eng 8:532–542.

61. Nissen TL, Hamann CW, Kielland-Brandt MC, Nielsen J, Villadsen J. 2000. Anaerobic and aerobic batch cultivations of Saccharomyces cerevisiae mutants impaired in glycerol synthesis. Yeast 16:463–474.

62. Merico A, Sulo P, Piškur J, Compagno C. 2007. Fermentative lifestyle in yeasts belonging to the Saccharomyces complex. FEBS J 274:976–989.

63. Kiers J, Zeeman A-M, Luttik M, Thiele C, Castrillo JI, Steensma HY, Dijken JP Van, Pronk JT. 1998. Regulation of alcoholic fermentation in batch and chemostat cultures of Kluyveromyces lactis CBS 2359. Yeast 14:459–469.

64. Stasyk O. 2017. Methylotrophic Yeasts as Producers of Recombinant Proteins, p. 325–350. In Sibirny, AA (ed.), Biotechnology of Yeasts and Filamentous Fungi.Springer International Publishing, Cham.

65. Meyers A, Weiskittel TM, Dalhaimer P. 2017. Lipid Droplets: Formation to Breakdown. Lipids 52:465–475.

66. Casey GP, Magnus CA, Ingledew WM. 1984. High-Gravity Brewing: Effects of Nutrition on Yeast Composition, Fermentative Ability, and Alcohol Production. Appl Environ Microbiol 48:639–646.

67. Wikén T, Scheffers WA, Verhaar AJM. 1961. On the existence of a negative pasteur effect in yeasts classified in the genus Brettanomyces Kufferath et van Laer. Antonie Van Leeuwenhoek 27:401–433.

68. Custers MTJ. 1940. Onderzoekingen over het Gistgeslacht Brettanomyces. TU Delft.

69. Tiukova IA, Petterson ME, Tellgren-Roth C, Bunikis I, Eberhard T, Pettersson OV, Passoth V. 2013. Transcriptome of the Alternative Ethanol Production Strain Dekkera bruxellensis CBS 11270 in Sugar Limited, Low Oxygen Cultivation. PLoS One 8:2–8.

70. Shen X-X, Zhou X, Kominek J, Kurtzman CP, Hittinger CT, Rokas A. 2016. Reconstructing the Backbone of the Saccharomycotina Yeast Phylogeny Using Genome-Scale Data. G3 Genes, Genomes, Genet 6:3927–3939.

71. Juergens H, Niemeijer M, Jennings-Antipov LD, Mans R, Morel J, van Maris AJA, Pronk JT, Gardner TS. 2018. Evaluation of a novel cloud-based software platform for structured experiment design and linked data analytics. Sci Data 5:1–12.

72. Juergens H, Varela JA, de Vries ARG, Perli T, Gast VJM, Gyurchev NY, Rajkumar AS, Mans R, Pronk JT, Morrissey JP, Daran J-MG. 2018. Genome editing in Kluyveromyces and Ogataea yeasts using a broad-host-range Cas9/gRNA co-expression plasmid. FEMS Yeast Res 1–16.

73. Gao J, Gao N, Zhai X, Zhou YJ. 2021. Recombination machinery engineering for precise genome editing in methylotrophic yeast Ogataea polymorpha. iScience 24:102168.

74. Medina VG, Almering MJH, Van Maris AJA, Pronk JT. 2010. Elimination of glycerol production in anaerobic cultures of a Saccharomyces cerevisiae strain engineered to use acetic acid as an electron acceptor. Appl Environ Microbiol 76:190–195.

75. Guadalupe-Medina V, Wisselink HW, Luttik MA, De Hulster E, Daran JM, Pronk JT, Van Maris AJA. 2013. Carbon dioxide fixation by Calvin-Cycle enzymes improves ethanol yield in yeast. Biotechnol Biofuels 6.

76. Papapetridis I, Goudriaan M, Vázquez Vitali M, De Keijzer NA, Van Den Broek M, Van Maris AJA, Pronk JT. 2018. Optimizing anaerobic growth rate and fermentation kinetics in Saccharomyces cerevisiae strains expressing Calvin-cycle enzymes for improved ethanol yield. Biotechnol Biofuels 11:1–17.

77. Alimardani P, Régnacq M, Moreau-Vauzelle C, Ferreira T, Rossignol T, Blondin B, Bergès T. 2004. SUT1-promoted sterol uptake involves the ABC transporter Aus1 and the mannoprotein Dan1 whose synergistic action is sufficient for this process. Biochem J 381:195–202.

78. Shi NQ, Jeffries TW. 1998. Anaerobic growth and improved fermentation of Pichia stipitis bearing a URA1 gene from Saccharomyces cerevisiae. Appl Microbiol Biotechnol 50:339–345.

79. Gojković Z, Knecht W, Zameitat E, Warneboldt J, Coutelis JB, Pynyaha Y, Neuveglise C, Møller K, Löffler M, Piškur J. 2004. Horizontal gene transfer promoted evolution of the ability to propagate under anaerobic conditions in yeasts. Mol Genet Genomics 271:387–393.

80. Lõoke M, Kristjuhan K, Kristjuhan A. 2011. Extraction of genomic DNA from yeasts for PCR-based applications. Biotechniques 50:325–328.

81. Juergens H, Hakkaart XDV., Bras JE, Vente A, Wu L, Benjamin KR, Pronk JT, Daran-Lapujade P, Mans R. 2020. Contribution of Complex I NADH dehydrogenase to respiratory energy coupling in glucose-grown cultures of Ogataea parapolymorpha. Appl Environ Microbiol 86:1–18.

82. Krügel H, Fiedler G, Smith C, Baumberg S. 1993. Sequence and transcriptional analysis of the nourseothricin acetyltransferase-encoding gene nat1 from Streptomyces noursei. Gene 127:127–131.

83. Goldstein AL, McCusker JH. 1999. Three New Dominant Drug Resistance Cassettes for Gene Disruption in Saccharomyces cerevisiae. Yeast 15:1541–1553.

84. Janke C, Magiera MM, Rathfelder N, Taxis C, Reber S, Maekawa H, Moreno-Borchart A, Doenges G, Schwob E, Schiebel E, Knop M. 2004. A versatile toolbox for PCR-based tagging of yeast genes: new fluorescent proteins, more markers and promoter substitution cassettes. Yeast 21:947–962.

85. Norrander J, Kempe T, Messing J. 1983. Construction of improved M13 vectors using oligodeoxynucleotide-directed mutagenesis. Gene 26:101–106.

86. Güldener U, Heck S, Fiedler T, Beinhauer J, Hegemann JH. 1996. A new efficient gene disruption cassette for repeated use in budding yeast. Nucleic Acids Res 24:2519–2524.

87. Entian K-D, Kötter P. 2007. 25 Yeast Genetic Strain and Plasmid Collections, p. 629–666. In Methods in Microbiology.

88. Fairhead C, Llorente B, Denis F, Soler M, Dujon B. 1996. New vectors for combinatorial deletions in yeast chromosomes and for gap-repair cloning using ‘split-marker’ recombination. Yeast 12:1439–1457.

89. Verduyn C, Postma E, Scheffers WA, van Dijken JP. 1990. Physiology of Saccharomyces cerevisiae in anaerobic glucose-limited chemostat cultures. J Gen Microbiol 136:395–403.

90. Luttik MAH, Kötter P, Salomons FA, van der Klei IJ, van Dijken JP, Pronk JT. 2000. The Saccharomyces cerevisiae ICL2 gene encodes a mitochondrial 2-methylisocitrate lyase involved in propionyl-coenzyme a metabolism. J Bacteriol 182:7007–7013.

91. Mashego MR, van Gulik WM, Vinke JL, Heijnen JJ. 2003. Critical evaluation of sampling techniques for residual glucose determination in carbon-limited chemostat culture of Saccharomyces cerevisiae. Biotechnol Bioeng 83:395–399.

92. Lange HC, Heijnen JJ. 2001. Statistical reconciliation of the elemental and molecular biomass composition of Saccharomyces cerevisiae. Biotechnol Bioeng 75:334–344.

93. Piper MDW, Daran-Lapujade P, Bro C, Regenberg B, Knudsen S, Nielsen J, Pronk JT. 2002. Reproducibility of Oligonucleotide Microarray Transcriptome Analyses. J Biol Chem 277:37001–37008.

94. Grabherr MG, Haas BJ, Yassour M, Levin JZ, Thompson DA, Amit I, Adiconis X, Fan L, Raychowdhury R, Zeng Q, Chen Z, Mauceli E, Hacohen N, Gnirke A, Rhind N, Di Palma F, Birren BW, Nusbaum C, Lindblad-Toh K, Friedman N, Regev A. 2011. Full-length transcriptome assembly from RNA-Seq data without a reference genome. Nat Biotechnol 29:644–652.

95. Singh R, Lawal HM, Schilde C, Glöckner G, Barton GJ, Schaap P, Cole C. 2017. Improved annotation with de novo transcriptome assembly in four social amoeba species. BMC Genomics 18:120.

96. Palmer J, Stajich J. 2019. funannotate. 1.7.1.

97. Langmead B, Trapnell C, Pop M, Salzberg SL. 2009. Ultrafast and memory-efficient alignment of short DNA sequences to the human genome. Genome Biol 10:R25.

98. Li H, Handsaker B, Wysoker A, Fennell T, Ruan J, Homer N, Marth G, Abecasis G, Durbin R. 2009. The Sequence Alignment/Map format and SAMtools. Bioinformatics 25:2078–2079.

99. Liao Y, Smyth GK, Shi W. 2014. FeatureCounts: An efficient general purpose program for assigning sequence reads to genomic features. Bioinformatics 30:923–930.

100. McCarthy DJ, Chen Y, Smyth GK. 2012. Differential expression analysis of multifactor RNA-Seq experiments with respect to biological variation. Nucleic Acids Res 40:4288–4297.

101. Robinson MD, Oshlack A. 2010. A scaling normalization method for differential expression analysis of RNA-seq data. Genome Biol 11.

102. Jones P, Binns D, Chang HY, Fraser M, Li W, McAnulla C, McWilliam H, Maslen J, Mitchell A, Nuka G, Pesseat S, Quinn AF, Sangrador-Vegas A, Scheremetjew M, Yong SY, Lopez R, Hunter S. 2014. InterProScan 5: Genome-scale protein function classification. Bioinformatics 30:1236–1240.

103. Lechner M, Findeiß S, Steiner L, Marz M, Stadler PF, Prohaska SJ. 2011. Proteinortho: Detection of (Co-)orthologs in large-scale analysis. BMC Bioinformatics 12:124.

104. Gu Z, Eils R, Schlesner M. 2016. Complex heatmaps reveal patterns and correlations in multidimensional genomic data. Bioinformatics 32:2847–2849.

105. R Core Team. 2017. R: A Language and Environment for Statistical Computing. Vienna, Austria.

106. Camacho C, Coulouris G, Avagyan V, Ma N, Papadopoulos J, Bealer K, Madden TL. 2009. BLAST+: Architecture and applications. BMC Bioinformatics 10:1–9.

107. Steenwyk JL, Rokas A. 2019. Treehouse: A user-friendly application to obtain subtrees from large phylogenies. BMC Res Notes https://doi.org/10.1186/s13104-019-4577-5.

